# Lung-mimicking click alginate-dECM model of breast cancer lung metastasis reveals the role of ECM mechanics in tumor growth dynamics and genomic instability

**DOI:** 10.64898/2026.01.08.698349

**Authors:** Laura Fallert, Ane Urigoitia-Asua, Jose Miguel Pardo-Sanchez, Elena Arizaga-Batiz, Rosa M. Zubizarreta-Susperregui, Dorleta Jimenez de Aberasturi, Amaia Cipitria

## Abstract

Metastatic breast cancer (BC) is the main cause of cancer-related death in women. Accumulating evidence highlights the prominent changes within the lung metastatic niche, resulting in stiffening of the extracellular matrix (ECM). The prevailing concept of cancer evolution encompasses the acquisition of beneficial traits due to genomic instability, yet it remains elusive to what extend altered lung ECM mechanics feed into this. To investigate this, a tunable 3D bioengineered model, capturing the biophysical and biochemical characteristics of the BC metastatic lung niche is developed. Porcine derived lung decellularized ECM (dECM), combined with norbornene and tetrazine modified click-crosslinkable alginate, recapitulates composition and mechanics of healthy soft (3 kPa) and metastatic stiff (12 kPa) lung niches. Proof-of-concept evaluation using label-free optical microscopy revealed preferential ECM fiber orientation in human lung metastatic tissue and in stiff hydrogels. Encapsulation of MDA-MB-231, MCF7 and patient-derived triple negative BC cells demonstrated that stiffer matrices were associated with DNA damage, indicated by yH2AX and altered Lamin A/C. Pharmacological perturbation of FAK mediated mechanotransduction reversed this phenotype in stiff conditions. These findings underscore the critical role of tissue mechanics in regulating BC metastasis and demonstrate the utility of the herein developed tunable, physiologically relevant platform for patient-based models.

**Table of Content:** 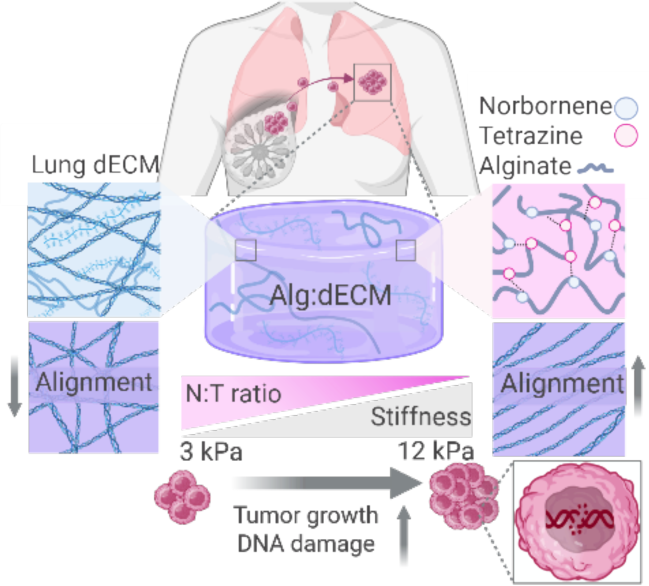

A bioengineered 3D model recreates mechanical and biochemical features of healthy and metastatic lung tissue. Metastatic-like stiff environments promote extracellular matrix organization, increased BC cluster growth and DNA damage, in both cell lines and patient-derived BC cells. These findings reveal how altered tissue mechanics may contribute to breast cancer progression and provide a platform for patient-derived models.

## 1. Introduction

Breast cancer (BC) remains an increasing global burden, being the leading cause of cancer related mortality in women world-wide.^[1]^ The ever-increasing mortality rates are primarily driven by the metastatic outgrowth at distant organs, including lungs, bones, liver and brain, with lungs comprising the second most common site.^[2]^ While metastatic onset varies based on subtype, curative treatment strategies remain challenging once patients display clinically detectable metastasis, resulting in poor prognosis.^[3]^ Despite the advances over the past decades, an unmet need remains to gain key insights into drivers of metastatic progression within the lungs, as well as appropriate models, reliably recapitulating BC behavior and predicting drug responses.

Considering that BC cells originate from lobular and ductal regions of the mammary glands, the lungs pose a biochemically and biophysically distinct microenvironment to metastasized BC cells. This becomes specifically relevant within the framework of the structure-function paradigm, where the surrounding tissue impinges on cellular behavior. While the local extracellular matrix (ECM) substantially contributes to both, physical as well as biochemical properties of a given tissue, resident cells probe such ECM derived cues in order to guide adhesion, proliferation, migration and differentiation.^[4]^ This paradigm however is challenged during tumorigenesis, which is accompanied by extensive remodeling of the surrounding ECM. In the context of BC lung metastasis, numerous studies reported elevated ECM deposition in the metastatic niche, even prior to BC cell arrival, establishing a fibrotic environment.^[5–7]^ As a consequence, tissue stiffness increases from 1.5-5 kPa in healthy lung tissue, up to 10-15 kPa in cancerous lung tissue.^[8–10]^ In addition to altered tissue mechanics, tumors also display prominent changes in tissue microarchitecture, characterized by pronounced ECM fiber alignment.^[11–13]^ It is well documented that increased ECM stiffness drives primary BC expansion, aggressiveness and progression, with accumulating evidence unraveling the mechanotransduction-driven proliferation, invasion and differentiation. ^[14–18]^ Whether this translates to metastatic BC cells within the lungs, exposed to a substantially distinct scale of tissue mechanics compared to their primary niche (0.92-0.96 kPa), has yet to be explored. ^[19]^ While multiple lines of in vivo work have shown that a fibrotic environment induces proliferative metastatic lung lesions, this has been primarily investigated in the context of fibrosis-associated changes in biochemical ECM composition rather than mechanics.^[20–22]^ Thus, an unmet need remains to understand direct and fundamental effects of changes in lung ECM mechanics on metastatic BC cell progression.^[23]^

Mechanotransduction occurs as cells integrate ECM derived mechanical stimuli from the cytoplasmic membrane, through the cytoskeleton, down to the cell nucleus. The nucleus is a mechanosensitive organelle, while also embodying a celĺs genomic storage unit. ^[24]^ Here, the nuclear envelope, composed of an inner and outer nuclear membrane and the nuclear lamina, comprises the nuclear-cytoplasmic interface, which communicates mechanical cues between cytoskeleton and nuclear interior. In particular the nuclear lamina, holds a pivotal role in maintaining nuclear integrity, and hence genomic stability, in response to mechanical cues.^[25,26]^ In fact, proteomic profiling of distinct tissues revealed that lamin A levels scale with tissue stiffness and collagen abundance, highlighting the importance of protecting nuclear content from external mechanical stress.^[27]^ Further supporting this notion, meta analyses revealed that genomic variation across solid cancers scale with the stiffness of the origin tissue.^[28]^ Likewise, chromosome loss and gains have been correlated with collagen abundance, and in vitro work has provided direct evidence of stiffness-induced chromosome loss. ^[29]^ Together, this argues for a central role of tissue mechanics in genomic integrity. Granted that genomic instability lies at the core of cancer evolution, to what extend stiffening of the BC metastatic lung niche contributes to this has yet to be determined.

Addressing such knowledge gaps requires sophisticated 3D models that capture multiple relevant aspects of the native target niche including mechanics, biochemical composition and microarchitecture. To date the leading biomaterials, specifically for modeling lung, remain single component inert or bioactive polymers such as i.e. alginate (Alg) and agarose or collagen, fibronectin and hyaluronic acid (HA), respectively.^[29–34]^ Alternatively, to yield higher degrees of mechanical tunability, UV-light mediated photo-crosslinkable semi-synthetic materials have been employed, such as gelatin methacrylate (GelMA), HA methacrylate (HAMA) or purely synthetic materials such as polyethylene glycol (PEG).^[35–40]^ Opposed to photo-crosslinkable systems, polymer systems with functional groups engaging in bio-orthogonal self-crosslinking click chemistry, such as norbornene (N) and tetrazine (T) modified Alg, provide excellent spatiotemporal control and dynamic tunability, at lower risk of potential cytotoxicity.^[41,42]^ Nonetheless, these materials fall short in capturing the niche-specific biochemical cues of lung tissue. While Matrigel remains a gold standard in organoid culture, owing to its biochemical complexity, it still fails to provide a physiologically appropriate microenvironment, due to its poor mechanical properties and its origin being murine sarcoma.^[43]^

More recently, organ derived decellularized ECM (dECM) has gained increasing popularity due to its capacity to provide a proteomic profile similar to the target tissue, making it a superior alternative to Matrigel. ^[44–46]^ However, a primary constraint remains a trade-off between ease of use at lower concentrations and mechanical properties, which not only fall far below lung tissue mechanics but also cannot be tuned independently of ligand density.^[47–49]^ Thus, achieving the desired tissue characteristics requires synergistic combinations of biomaterials which deliver mechanical tunability as well as the appropriate biochemical profile. Limited platforms incorporating lung mechanical as well as bioactive cues have emerged, serving as the blueprint for advanced bio-engineered models. In this regard, PEG-based systems have captured healthy lung tissue mechanics and composition by incorporating a selected set of prevalent synthetic peptides or corresponding sequences, evident in human lung.^[31,39,40]^ While these provide controlled and reproducible approaches, an unmet need remains to fully capture the native niche, including tissue mechanics, composition and microarchitecture.

Here we aim to improve the understanding of BC lung metastasis in the context of a physiologically relevant physical and biochemical microenvironment. We hypothesize that a stiffer lung ECM drives growth and genomic instability of BC cells. For this purpose, we developed a biomimetic 3D in vitro model based on porcine lung dECM and click alginate, with tunable tissue mechanics and fibrous architecture to recapitulate the native lung metastatic niche on a physical and biochemical level. We leverage this model to decipher BC cell cycle and growth dynamics, as well as genomic instability in both cell lines and patient-derived BC cells. These findings reveal how altered tissue mechanics may contribute to breast cancer progression and provide a platform for patient-derived models.

## 2. Results

### 2.1. Proof-of concept characterization of patient BC lung metastatic ECM architecture

#### 2.1.1. Label-free imaging reveals preferential ECM fiber orientation in patient BC lung metastatic tissue

First, to determine characteristics of the microarchitecture within the lung metastatic niche in patients, we performed advanced label-free imaging on unstained tissue sections of paraffin embedded lung metastatic tissue samples of BC patients. Second harmonic generation (SHG) and two photon excitation fluorescence (TPEF) based signals allow for label-free visualization of collagen and elastin fibers, respectively. ^[50,51]^ Upon consultation with a trained pathologist, the tumor cell enriched regions based on corresponding hematoxylin and eosin (H&E) staining were determined (Supplementary Figure S1A). Given that specifically ECM fiber alignment has been a reported feature in the context of BC and lung cancer, we performed CTfire based analysis to quantify angular distributions of collagen and elastin fibers (Figure 1B-C).^[52,53]^ This revealed that both ECM components exhibited preferential, non-random angular distributions, rather than uniformly distributed orientations. Individual samples showed distinct peaks in fiber orientation, indicating preferentially oriented fiber populations, while position and breadth of distributions varied across patients.

**Figure 1.**
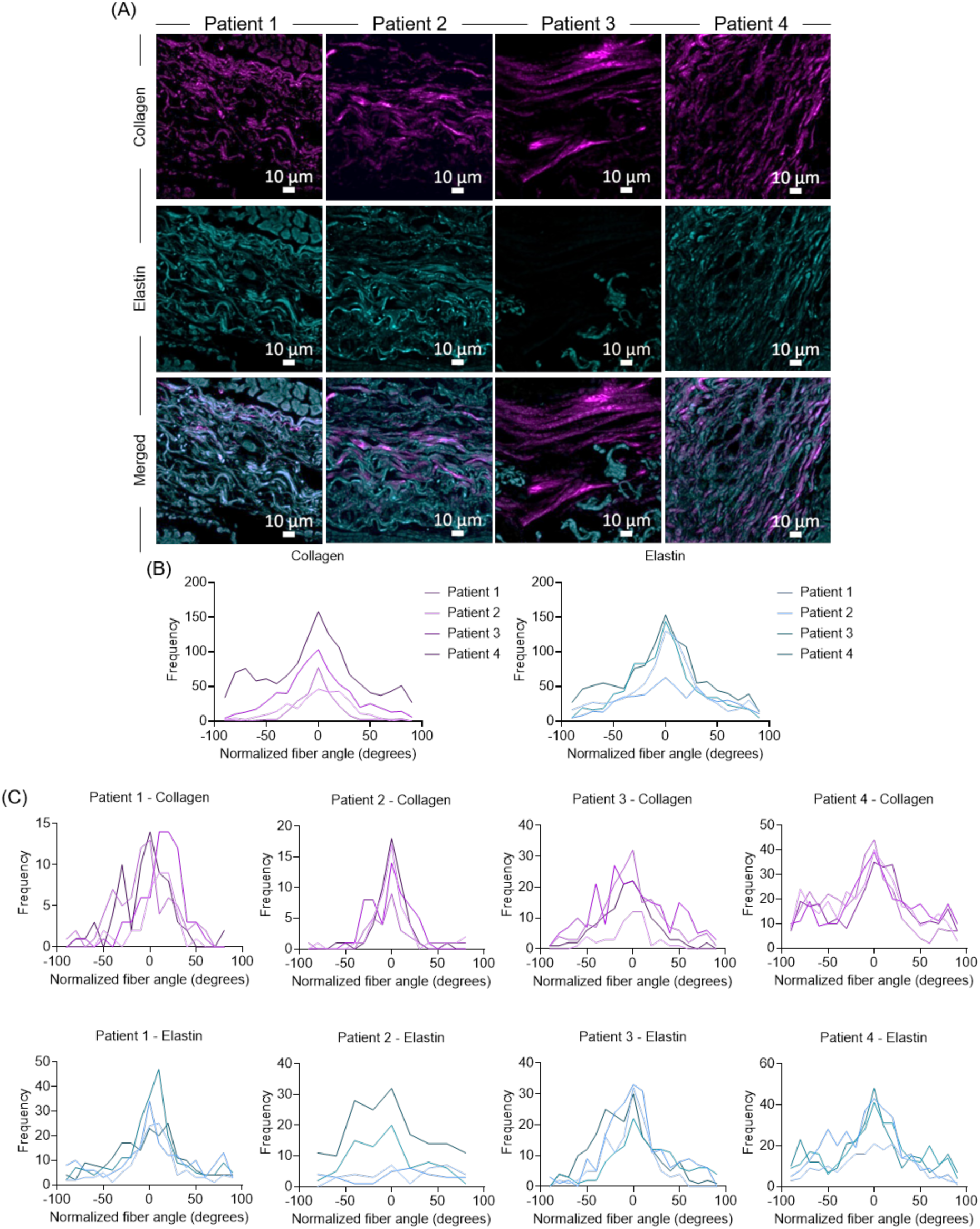
Collagen and elastin fiber distributions in patient BC metastatic lung samples. A) Second harmonic generation (collagen, magenta) and two photon excitation fluorescence (elastin, cyan) images of patient BC lung metastatic paraffin tissue sections. B) Fiber angle distribution of normalized collagen and elastin fibers across patients, where each line represents the sum of each region and C) within different regions in each patient, where each line represents individual regions per patient, respectively. B) n = 4 donors and C) n = 4 regions/donor.

### 2.2. Development and characterization of tunable lung mimicking ECM

#### 2.2.1. Decellularized porcine lung ECM retains core lung ECM constituents evident in human lung

Next, we sought to recapitulate the biochemical composition of the lung niche in vitro, relying on porcine lung derived dECM. As human lung tissue of healthy donors is scarce, opting for porcine-derived lung tissue is an attractive alternative source to obtain organ specific dECM. The primary objectives of the decellularization process are the complete removal of porcine cells and minimizing any residual DNA, while maintaining ECM constituents. Once collected, lungs were dissected, bronchi and bronchioles were removed, and the remaining tissue was aliquoted and stored at −80°C until decellularization. Upon decellularization of dissected lung tissue, histological assessment revealed that elastin and collagen were well preserved (Figure 2A). H&E staining on the other hand confirmed the removal of cellular constituents, which we further validated via DNA quantification, showing significantly lower DNA levels in decellularized tissue compared to native tissue (Figure 2A-B).

**Figure 2.**
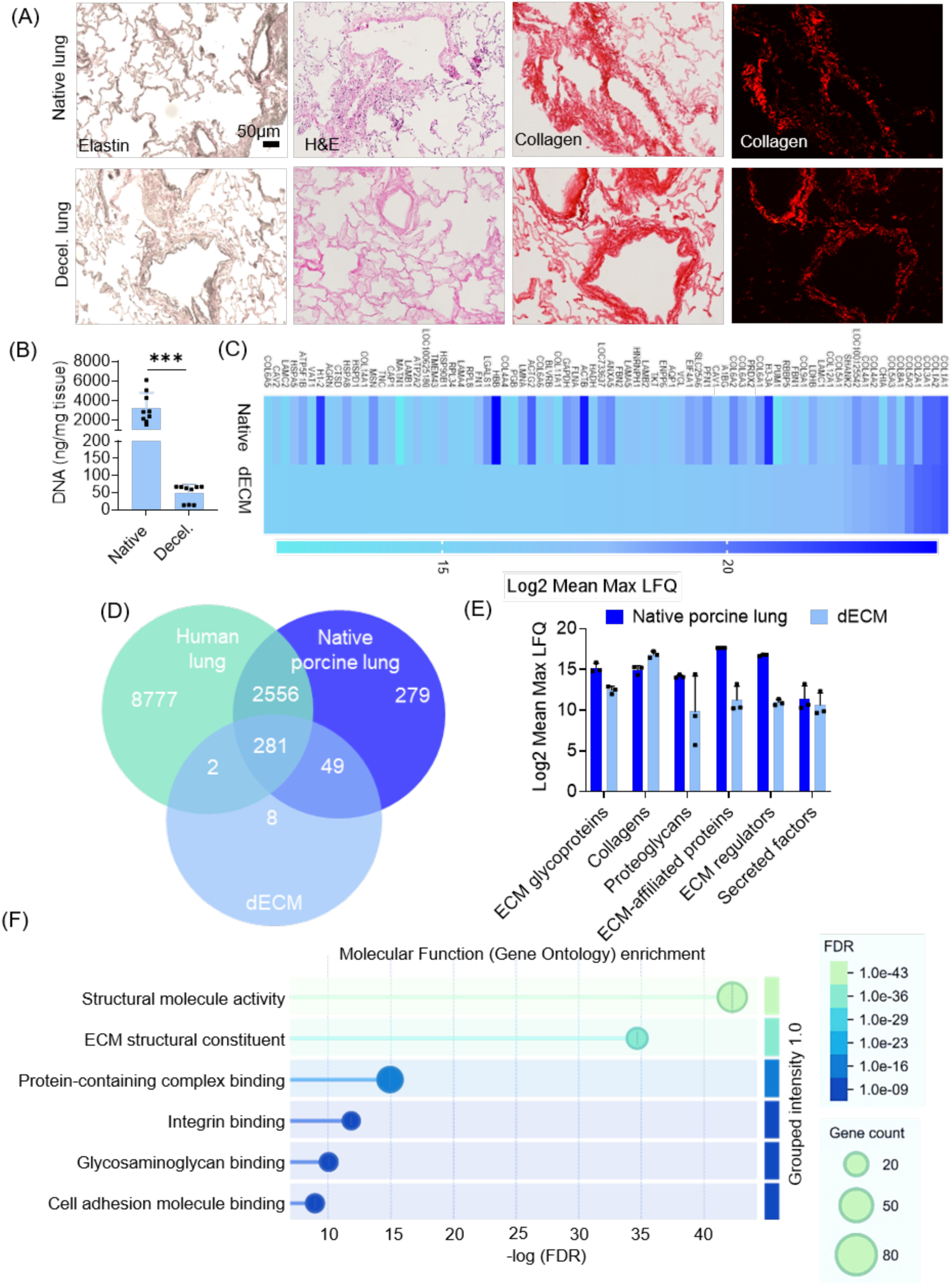
Lung dECM characterization. A) Representative bright field images of Van Gieson (Elastin), H&E, Picrosirius red (collagen, brightfield left; polarized light, right) stained native and decellularized (decel.) paraffin embedded porcine lung tissue. B) DNA quantification of native and decel. porcine lung tissue. C) Heatmap of the 80 most abundant genes coding for proteins identified in porcine lung dECM compared to native porcine lung tissue. D) Venn diagram depicting overlap across total human lung, native porcine lung and porcine lung dECM proteome. E) Graph displaying abundance of matrisomal proteins in native porcine lung in comparison to porcine lung dECM. F) Gene ontology analysis (STRING) of human orthologs evident in porcine lung dECM shows significant enrichment in terms involved in ECM-related functions. Terms were filtered based on a false discovery rate (FDR) of >0.05 and the graph shows log-transformed FDR values, with node size representing the number of genes associated with each gene ontology term, and node color intensity reflecting the −log(FDR). n = 3 donors. Bar graphs show mean with standard deviation. Student’s t-test. ***p≤0.001

To gain a comprehensive overview of the protein profile of native lung tissue and the resulting dECM, proteomic analysis was performed. This revealed that several core ECM proteins (collagens, laminins, fibronectins) retain abundances comparable to the native porcine lung tissue (Figure 2C). Of note, multiple of the genes coding for identified proteins in dECM (i.e. TNC, CAV1, FN1, FBN2 COL181, COL7A1) have been relevant in the context of BC lung metastasis. ^[12,39,54,55]^ To then determine cross-species differences, we assessed a publicly available human lung proteome dataset validating a well conserved cross-species protein composition, with >80 % of porcine native lung as well as dECM proteome overlapping with the human lung proteome (Figure 2D).^[56]^ Mapping the identified proteins to the matrisome database, a curated catalogue of human ECM constituents, further revealed that specifically matrisomal proteins are highly preserved upon decellularization (Figure 2E).^[57]^ In addition, we mapped a public Matrigel proteomic dataset to the matrisome data base and discovered that only 10 % of detected genes in Matrigel correspond to matrisome components, whereas 25 % of our porcine lung dECM proteins are annotated as matrisome components (Supplementary Figure S2A).^[45]^ This can be explained by the fact that Matrigel is tumor derived and thus tends to be very rich in growth factors rather than ECM proteins. We further leveraged this data to perform functional enrichment analysis, revealing a significant over-representation of terms with ECM-related molecular functions in porcine dECM (Figure 2F). Subsequently, we assessed a public human lung ECM-isolate proteomic dataset, to determine and compare reactome pathways across porcine lung dECM and Matrigel.^[58]^ As expected, this revealed prominent associations of both, porcine lung dECM and human lung ECM proteins with ECM related pathways including ECM organization, collagen formation, degradation, trimerization, assembly and biosynthesis, as well as integrin cell surface interactions and ECM proteoglycans (Supplementary Figure S2B-C). While porcine lung dECM and human lung ECM pathway enrichment displayed a high degree of resemblance, Matrigel displayed primarily metabolic and translation related pathways, rather than ECM related (Supplementary Figure S2D). This aligns with our prior observation of limited matrisomal annotations evident in Matrigel.

Taken together, the established porcine derived lung dECM provides a physiologically informed and compositionally diverse organ-specific extracellular environment, reflecting the native target tissue niche, and thus, making it a suitable tool to introduce biochemical complexity in a 3D bioengineered lung metastasis model.

#### 2.2.2. Click Alg-dECM captures essential features of the lung metastatic niche

Lung dECM mechanical properties are determined by the amount of physical collagen crosslinks and therefore depend on the dECM concentration. This impedes the separation of effects driven by mechanical cues and effects driven by ligand density. Moreover as already noted, lung dECM in itself exhibits poor mechanical properties falling below the moduli of native lung.^[47,49]^ To circumvent this and to decouple ligand density from mechanical properties, we implement click N-T-Alg, providing a high degree of tunability by changing N:T ratios.^[41,42]^ While mechanical properties reported for native lung tissue are derived from different measurement modalities, therefore cannot be considered directly equivalent to rheologically-determined hydrogel properties, these values were used as a reference range to guide hydrogel design. Upon synthesizing low and high molecular weight (LMW & HMW) N-Alg and T-Alg in combination with lung dECM (Alg-dECM) yielded an entirely self-crosslinkable hydrogel, owing to the bio-orthogonal, covalent inverse-electron-demand Diels-Alder crosslinks between N-T-Alg and the physical collagen crosslinks of dECM. Moreover, as expected Alg-dECM hydrogels display a high degree of tunability, with formulations based on LMW N-T-Alg falling within the range of healthy lung tissue of 1.5-5 kPa (Supplementary Figure S3C).^[8,9]^. However, we opted to rely on HMW N-T-Alg based formulations to achieve both moduli of healthy as well as cancerous lung tissue (10-15 kPa), from here on referred to as soft and stiff, respectively.^[59]^ More specifically, the final formulations were based on a constant composition of 2.5 % w/v HMW Alg, with or without 0.5 % w/v dECM, and varying crosslinking degree with N:T 4.0 yielding soft (Alg-dECM 3.62 ± 0.91 kPa; Alg, 3.187 ± 388 kPa) and a N:T 0.5 yielding stiff (Alg:dECM 13.92 ± 1.50 kPa; Alg 11.306 ± 1.856 kPa) (Figure 3C-E). HMW Alg only, without dECM serves as a reference material, displaying matched mechanical properties at equivalent N:T ratios (Figure 3C-E).

**Figure 3.**
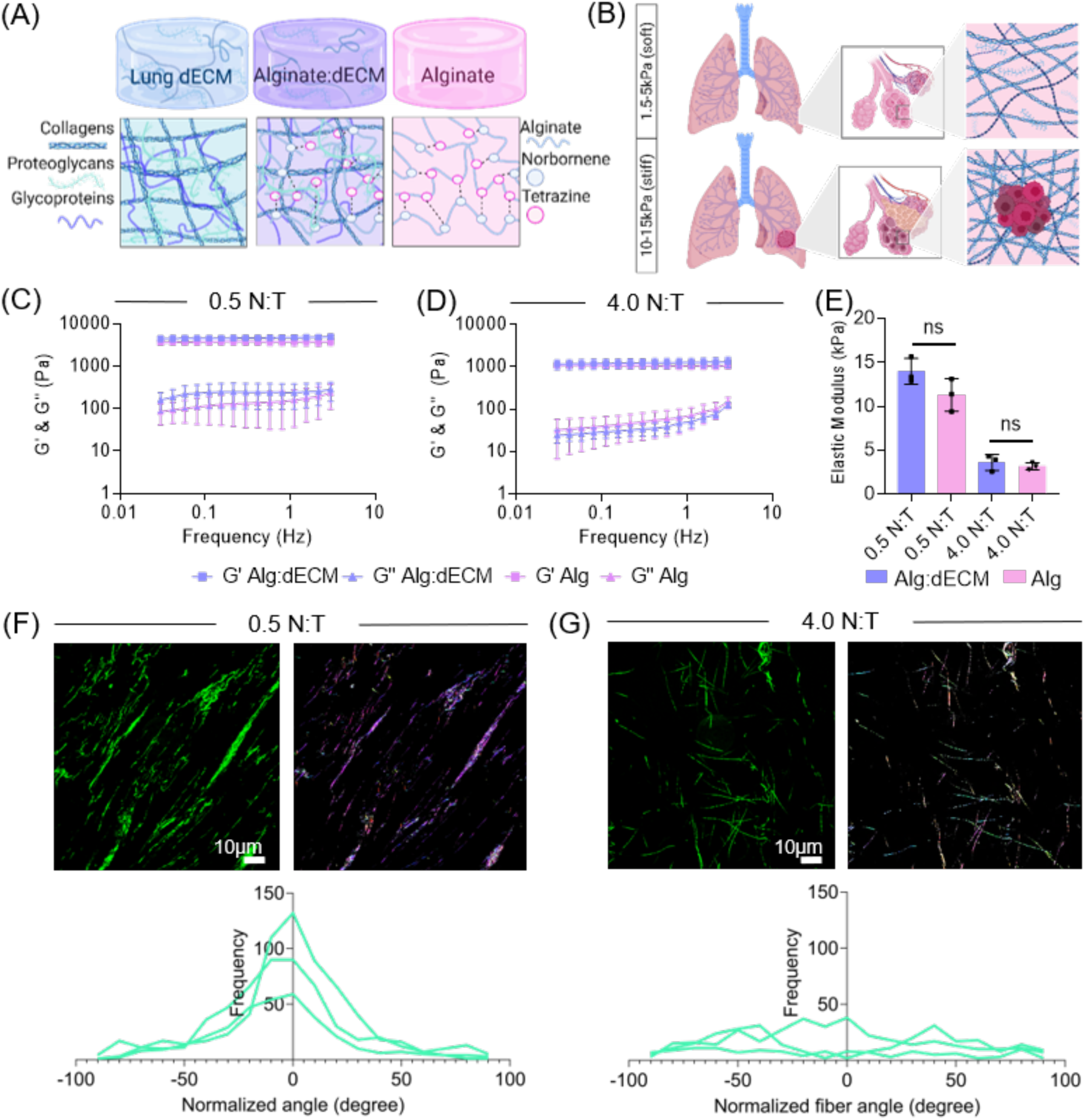
Characterization of click Alg-dECM based healthy and metastatic lung niche. A) Schematic representation of individual and hybrid polymer networks. B) Schematic representation of healthy and metastatic lungs with corresponding reported mechanical properties.^[8–10]^ Frequency sweeps of 2.5 % w/v HMW Alg with (purple) and without (pink) 0.5 % dECM at C) N:T 0.5 and D) N:T 4.0 with corresponding storage (G’) and loss (G’’) moduli. E) Elastic moduli of 2.5 % HMW Alg-dECM and Alg at N:T 0.5 and at N:T 4.0. F) Representative confocal reflectance images, orientation maps and corresponding fiber angle frequency distributions showing preferential collagen fiber orientation in stiff (0.5 N:T) compared to G) random fiber organization in soft (4.0 N:T), both with constant 2.5 % w/v HMW Alg with 0.5 % w/v dECM composition. n = 3 gels per condition. Graphs in C, D and E show mean with standard deviation. Graphs in F and G display distribution of normalized fiber angles. Mann-Whitney U test.

Furthermore, as demonstrated in Figure 1, the herein assessed metastatic lung tissue architecture displays preferential ECM fiber orientation. We used confocal reflectance imaging in order to assess collagen fiber architecture within our stiff vs. soft Alg-dECM hydrogels, mimicking metastatic vs. healthy lung tissue, respectively. This technique does not require sample fixation or staining and thus allows the visualization of the native state of collagen fibers and revealed a preferential collagen fiber orientation in stiff Alg-dECM compared to a random fiber orientation in soft Alg-dECM (Figure 3F-G).

Together, the above-described hydrogel formulations yield a lung mimicking ECM that captures the biochemical complexity, mechanical properties as well as architectural features of the target tissue, providing a physiologically relevant platform to study BC lung metastasis.

### 2.3 Characterization of BC cell behavior in mechanically tuned lung mimicking ECM

#### 2.3.1 BC cells display dynamic cell cycle patterns in mechanically tuned ECM

Initially, we aimed to verify the biocompatibility of the final formulations to exclude any potential dECM associated toxicity. While adequately decellularized and sufficiently washed dECM is generally considered biocompatible, the decellularization process involves several harsh reagents, whose residues could compromise cell viability. ^[60,61]^ For this purpose, viability tests were performed across all conditions with MCF7, representative of luminal ER^+^ BC, and MDA-MB-231 (MDA), representing basal like triple negative BC (TNBC). As both cell lines are derived from pleural effusions, they are a suitable tool to model lung metastasis across two distinct BC subtypes, with MDA displaying a highly metastatic and invasive mesenchymal profile, and MCF7 displaying a less invasive, epithelial profile. This demonstrated the biocompatibility of the final formulations across both BC cell subtypes and prompted further investigation of BC cell cycle dynamics as a function of ECM properties (Supplementary Figure S5). For this purpose, we employed genetically modified fluorescent ubiquitination-based cell cycle indicator 2 (FUCCI2) modified MDA and MCF7 cells, previously established in our group.^[62–64]^ Intermitted imaging of cell cycle distribution over culture periods of 2 weeks demonstrated proportional distribution of G1 and G2 phase cells across encapsulated MDA cells (Figure 4A,C-D). MCF7 cells on the other hand displayed G1 enriched clusters which is to be expected and can be explained by their well-known slower population doubling time opposed to MDA cells. (Figure 4B,E-F). Regarding overall growth across ECM formulations, MCF7 tended to display more growth in soft matrices compared to stiff at day 7, which equalizes at day 14 (Figure 4E, F), while MDA displayed comparable growth at day 7 but a tendency of more growth in stiff matrices on day 14 (Figure 4C, D). Together this highlights the compatibility of this platform with distinct BC subtypes and their dynamic growth patterns as a function of time.

**Figure 4.**
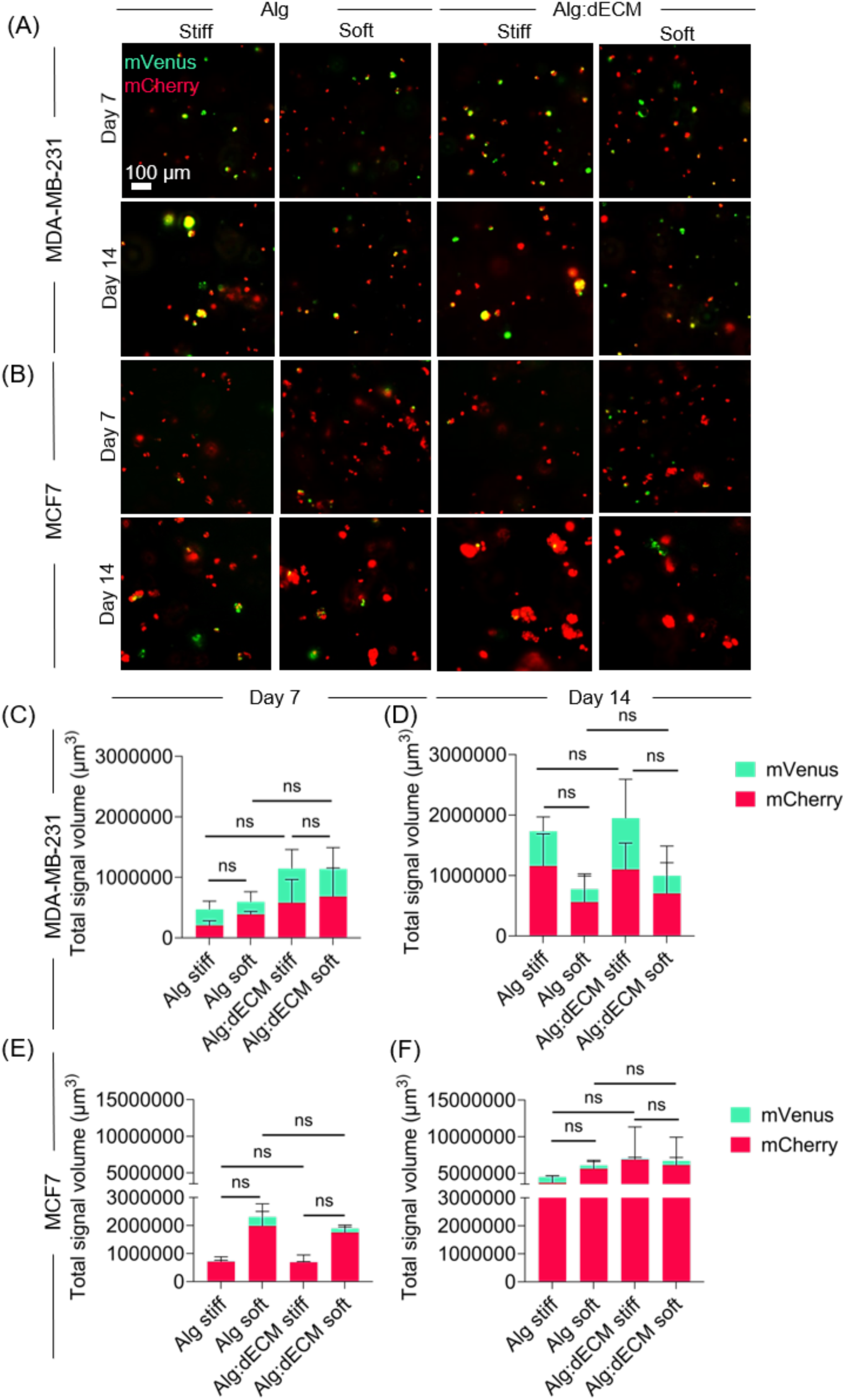
Cell cycle dynamics within mechanically tuned hydrogels. A) Representative images of Z-stack projections (200 µm) of FUCCI2 modified MDA-MB-231 and B) MCF7 cells over 14 days. C-D) Corresponding graphs displaying quantified total mCherry and mVenus signal volume of MDA-MB-231 and E-F) MCF7, across indicated hydrogel formulations over time. n = 3 gels per condition. Bar graphs show mean with standard deviation. Mann-Whitney U test.

#### 2.3.2 ECM stiffness promotes BC cluster growth and determines cluster morphology

Next, we aimed to determine changes across cluster morphologies as a function of ECM properties. For this purpose, cytoskeletal (Phalloidin) and nuclei (Dapi) staining of MDA and MCF7 cells were performed after a culture period of 21 days. This revealed significantly larger cluster volumes in stiff ECMs compared to soft ECMs across both cell lines (Figure 5 A,B,C,F). Quantification of additional morphological features revealed that stiff ECMs yielded more compact clusters, reflected in a significantly higher degree of sphericity and solidity compared to soft ECMs (Figure 5 D-E, G-H). MCF7 displayed generally larger clusters compared to MDA, reflecting their epithelial, adherent phenotype, whereas MDA clusters were smaller, reflective of their mesenchymal phenotype. Assessing the effect of the material composition, we observe significantly larger MDA clusters in Alg-dECM compared to Alg only (Figure 5C), suggesting a higher responsiveness of MDA, but not MCF7 cells, to the addition of lung dECM. Together, these results indicate a consistent effect of ECM stiffness, promoting BC cluster growth across distinct BC subtypes, and suggest a sensitivity of MDA cells towards lung derived factors.

**Figure 5.**
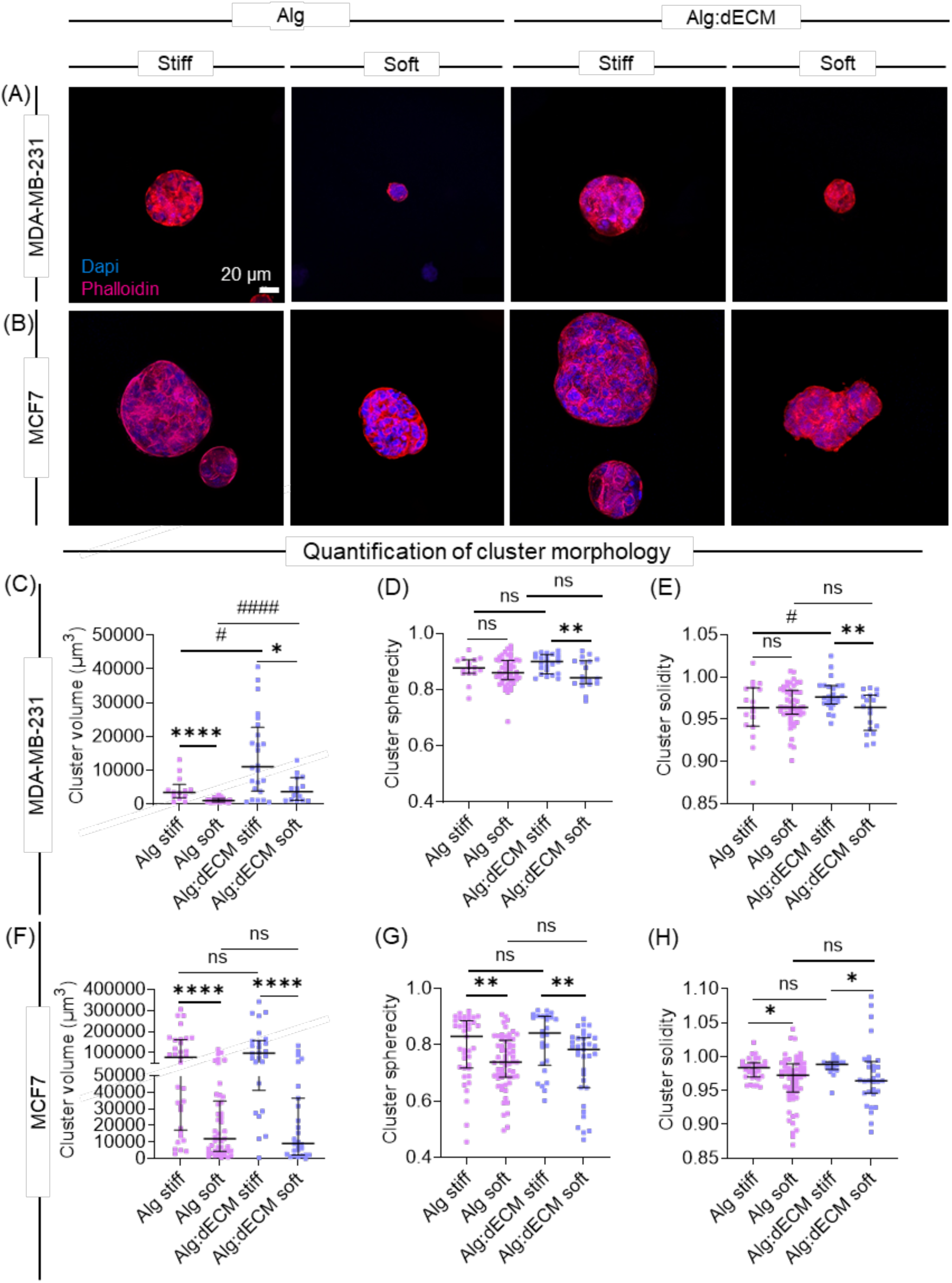
Matrix stiffness drives BC cluster growth and determines morphology. A) Representative images of Z-stack projections (30 µm) of Dapi (blue) and Phalloidin (red) stained MDA-MB-231 and B) MCF7 clusters after 21 days of encapsulated culture. C-E) Graphs showing quantification of morphological features of MDA-MB-231 and F-H) MCF7 cluster analysis. n =3 gels per condition. Graphs show median with interquartile range. Student’s t-tests were performed for normally distributed data and Mann-Whitney U tests were performed non-normally distributed data. Differences between stiffness is indicated by * and differences between Alg and Alg-dECM are indicated by #. */#p≤0.05, ** p≤0.01, ****/####p≤0.0001

#### 2.3.3 Stiff ECM is associated with genomic instability

Most studies investigating mechanobiology in the context of BC have focused on transient phenomena such as changes in morphology, transcription patterns and signaling pathway activation. However, to investigate how cancer associated changes in ECM mechanics ultimately contribute to changes driving cancer evolution, we assessed genomic integrity of BC cells in soft and stiff matrices. Given that lamin A/C serves as a nuclear mechanosensory and plays a pivotal role in preserving nuclear and hence genomic integrity, we performed nuclear lamin A/C stainings. Co-stainings with γH2AX, a DNA double strand break marker acting as a proxy for genomic instability, were performed to determine DNA damage patterns. After 14 days of culture in stiff and soft ECM, MDA cells displayed a reduction in lamin A/C levels in stiff conditions, which was accompanied by significantly elevated γH2AX levels, suggesting a stiffness-induced DNA damage (Figure 6A,C). MCF7 on the other hand, displayed an increase in lamin A/C levels in stiffer ECM (Figure 6B,E). However, this increase seemed insufficient in preserving genomic integrity, as MCF7 cells also displayed higher DNA damage in stiff compared to soft ECM (Figure 6B,F).

**Figure 6.**
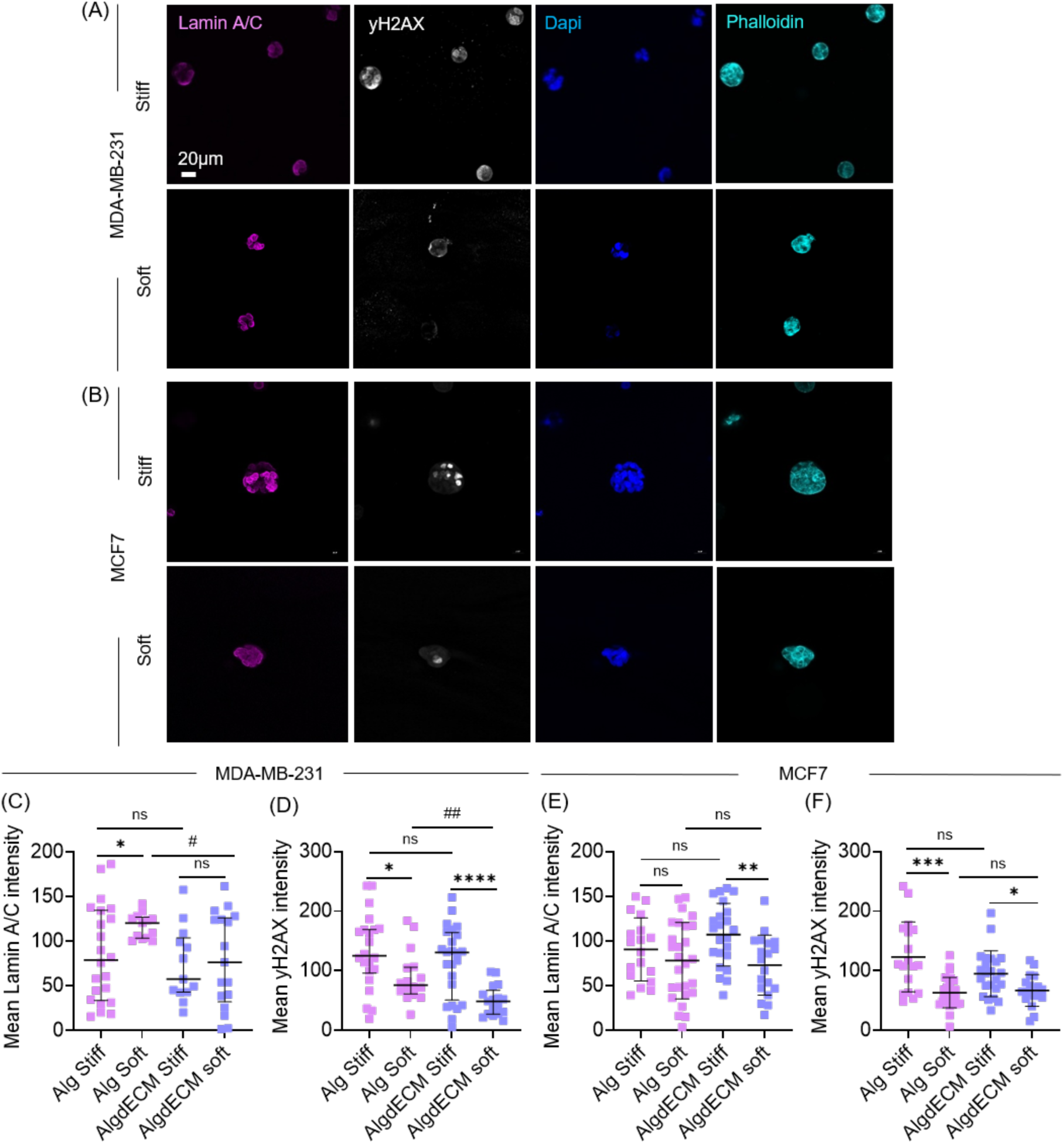
Matrix stiffness drives DNA damage in BC cells. A) Representative Z-stack projections (30 µm) of MDA and B) MCF7 cells cultured for 14 days in soft and stiff alginate-dECM, stained for nuclear lamin A/C, DNA damage marker γH2AX, Dapi and Phalloidin. C, D) Graphs showing quantified lamin A/C and γH2AX levels in MDA and E, F) MCF7 cells.. n = 3 gels per condition. Graphs show median with interquartile range. Student’s t-tests were performed for normally distributed data and Mann-Whitney U tests were performed for non-normally distributed data. Differences between stiffness is indicated by * and differences between Alg and Alg-dECM are indicated by #. */#p≥0.05, **/## p≥0.01,*** p≥0.001,****p≥0.0001.

Taken together, this data indicates the potential of stiffer ECM contributing to changes in genomic integrity in BC cells across distinct molecular subtypes, further eluding on the role of tissue mechanics on metastatic progression.

### 2.4 Model validation using patient-derived TNBC cells

#### 2.4.1 Compatibility of Alg-dECM with patient-derived triple negative BC cells

Finally, to assess the translational value of our platform, patient-derived primary TNBC cells were obtained and encapsulated in all four hydrogel formulations. Given that TNBC displays high frequencies of lung metastasis we opted for this subtype. Viability was assessed after 1, 7 and 14 days, revealing high viability with no pronounced differences across formulations (Figure 7A-B).

**Figure 7.**
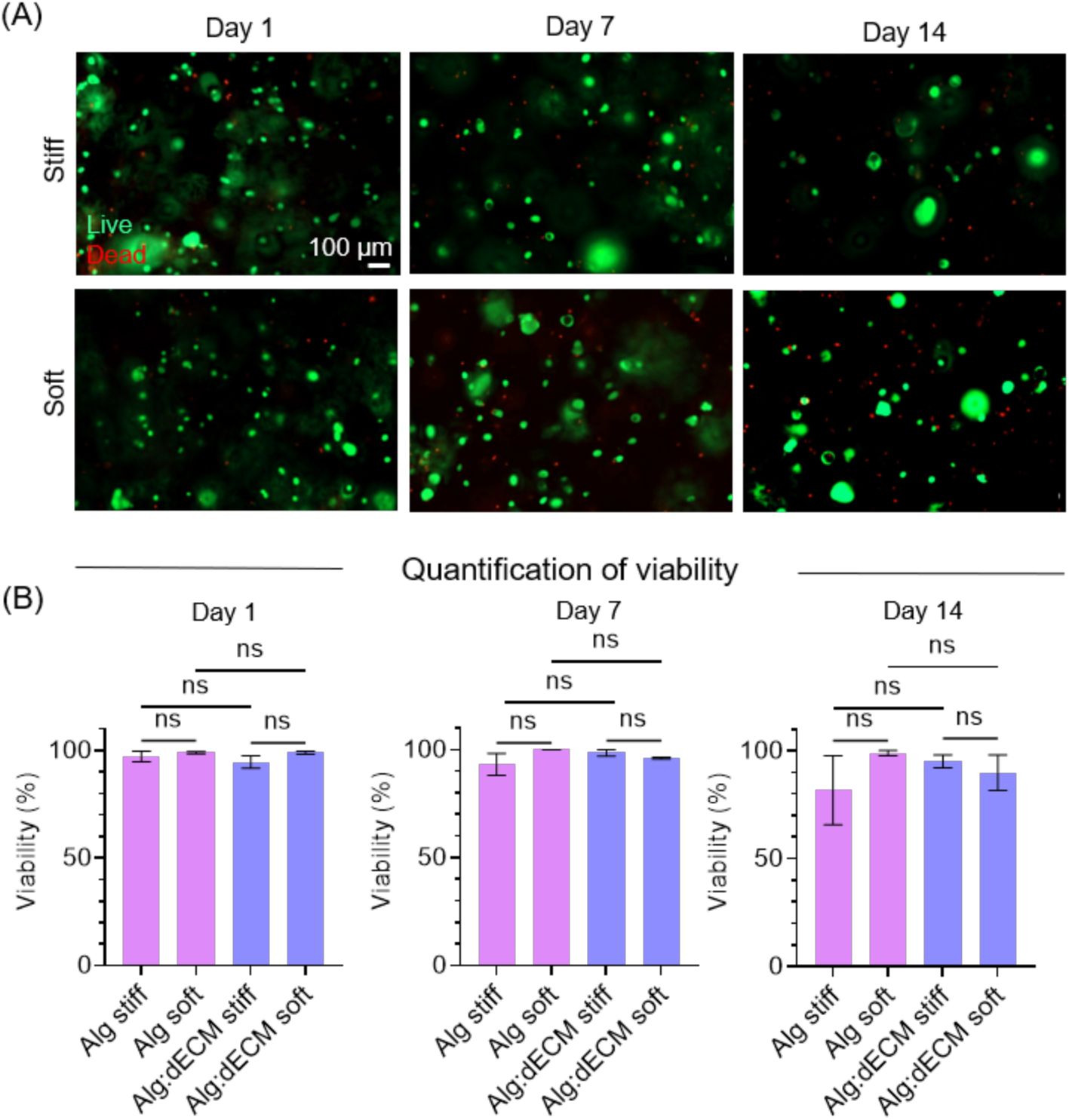
Mechanically tuned matrices provide a suitable platform for patient-derived TNBC cells. A) Viability assay with calcein-AM and ethidium homodimer-1 and representative images of Z-stack projection of patient-derived TNBC cells encapsulated in soft and stiff matrices, after 1,7 and 14 days of culture. B) Corresponding quantification of viability of patient-derived triple negative BC cells in indicated formulations. n = 3 gels per condition. Graphs show mean with standard deviation. Mann-Whitney U test.

#### 2.4.2 Focal adhesion kinase (FAK) inhibition reverts stiffness associated phenotype in patient-derived TNBC cells

To determine whether patient-derived cells exhibit similar stiffness-associated changes in genomic integrity and to functionally investigate the contribution of stiff matrices to elevated DNA damage, we targeted focal adhesion kinase (FAK), a central mediator of integrin dependent mechano-signaling.^[65]^ To this end, patient-derived TNBC cells cultured in stiff and soft Alg-dECM hydrogels were exposed to Defactinib, a clinically relevant FAK inhibitor (Figure 8).^[66]^ In vehicle-treated controls (0.1% v/v DMSO), stiff matrices were associated with significantly lower Lamin A/C levels and increased γH2AX levels compared with soft matrices, consistent with our previous observations in MDA-MB-231 cells, suggesting that matrix stiffness induces a conserved mechanical stress response in TNBC cells (Figure 8A-C). Pharmacological FAK inhibition, using up to 5 μM Defactinib, markedly altered this phenotype in stiff matrices, resulting in increased Lamin A/C levels and a substantial reduction in γH2AX (Figure 8A-C). In contrast, Lamin A/C and γH2AX levels were comparatively less affected by FAK inhibition in soft matrices (Figure 8A-C).

**Figure 8.**
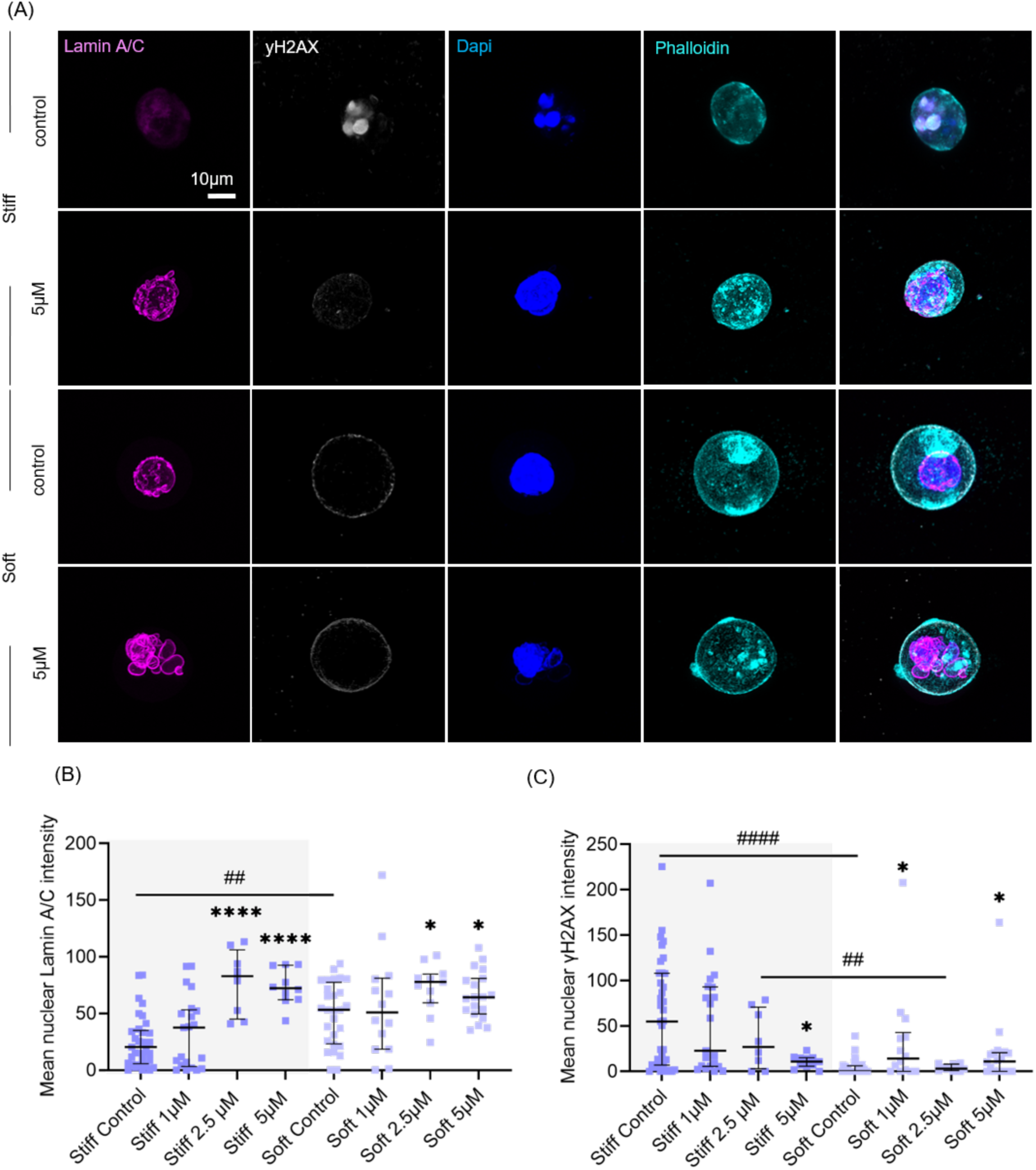
Matrix stiffness drives DNA damage in patient-derived TNBC cells, while FAK inhibition restores Lamin A/C and reduces DNA damage levels. A) Representative confocal images of Z-stack projections (30 µm) of patient-derived TNBC cells encapsulated in stiff and soft Alg-dECM for 14 days with and without Defactinib. B) Quantification of γH2AX and C) Lamin A/C levels at various Defactinib concentrations. n = 3 gels per condition. Graphs show medians with interquartile range. Student’s t-tests were performed for normally distributed data and Mann-Whitney U tests were performed for non-normally distributed data. Differences between concentrations are indicated by *, differences between stiffness are indicated by #.*p≥0.05, ##p≥0.01,****/####p≥0.0001

Collectively, this data indicates that FAK signaling contributes to stiffness-associated alterations in nuclear architecture and DNA damage, linking matrix derived mechanical forces to genomic stability in patient-derived TNBC cells.

## 4. Discussion

In the present study we lay down a patient-data driven 3D platform to recapitulate biophysical and biochemical aspects of the native metastatic lung niche. The combination of the herein established lung dECM with click N-T-Alg offers an entirely self-crosslinkable platform, with independent tunability of mechanics and bioactivity. Within this framework we demonstrated mechanical-driven growth and genomic instability, across distinct BC molecular subtypes. Owing to its compatibility not only with conventional BC cell lines but also with patient-derived TNBC cells, this platform holds potential as a personalized drug testing platform.

The above discussed data highlights the utility of tissue derived dECM as an attractive strategy to engineer physiologically relevant microenvironments. Our optimized protocol yields efficient removal of porcine cells and DNA traces. Proteomic data demonstrates that the final dECM not only retains matrisomal protein abundance, but also displays a high degree of overlap with the human lung proteome, offering an improved representation of the in vivo lung niche compared to single compound materials or Matrigel. This is in line with previous work highlighting the resemblance between human and porcine (decellularized) lung tissue.^[67]^ Importantly, the incorporation N-T-Alg provides a strategy addressing the mechanical mismatch between lung dECM and lung tissue.^[47–49]^ In addition, this approach improves the limited mechanical tunability, which is inherently tied to polymer concentration, impeding the distinction between ligand and stiffness-induced effects. We demonstrated that opting for different N:T ratios enables the decoupling of stiffness from ligand density. Another important aspect remains the more ethical sourcing of the starting material, as the collected lungs were a byproduct of standard surgical practice procedures within the local animal facility.

Moreover, the herein demonstrated collagen fiber architecture as a function of stiffness provides a simple methodology, omitting the need of external manipulation tools. The observed preferential collagen orientation in stiff Alg-dECM is likely due to the stiffness-related confinement imposed during collagen fibrillogenesis.^[68]^ With Alg being the dominant component, a denser polymer network in lower N:T ratios likely restricts isotropic growth of collagen fibrils, opposed to a looser network within higher N:T ratios. Importantly, as already mentioned, in clinical samples of primary BC it is well established that fiber alignment is not a trivial characteristic but rather holds prognostic value, further reflected in established tumor associated collagen signatures. ^[11,12,69]^ While previous work has indicated a similar tendency of collagen alignment within the lung metastatic niche in a mouse model, as well as in lung cancer patients, our data further supports the notion that this might also be a defining feature within human BC lung metastatic tissue. ^[53,55]^

Leveraging this platform, we revealed dynamic growth patterns, with MCF7 displaying G1 enriched clusters and MDA displaying a more proportional distribution, which reflects their distinct population doubling times. While we initially observed more growth of MCF7 cells in soft ECMs, this pattern was lost, with more growth developing between day 7-14 in stiff matrices. Our phalloidin based cluster morphology analysis on day 21 further revealed the development of larger and more spherical, solid clusters in stiff matrices of both, MCF7 as well as MDA cells. In line with this, in primary BC mimicking studies employing stiffness ranges from 0.1-5 kPa, a stiffness-induced increase in proliferation was observed. ^[14–16,18,70,71]^ However, previously MCF7 clusters encapsulated in matrices ranging from 2-17 kPa have been reported to display larger volumes in softer matrices but a higher density of nuclei in stiff matrices. ^[72,73]^ Such discrepancies are likely a result of distinct biomaterials employed, which despite displaying the same stiffness ranges, can differ substantially on multiple levels. For instance, it has been demonstrated that the presence and density of bioactive ligands (i.e. collagen), as well as stress relaxation time differentially impact BC cell behavior and cluster morphology at equivalent stiffness, impeding direct comparisons across studies.^[40,74,75]^

Regarding effects of dECM on BC cell behavior, we observe limited differences between Alg and Alg-dECM, likely due to the low dECM concentrations compared to Alg. The primary objective of this study was to harness the effect of tissue mechanics in a physiologically relevant setting and previous work has demonstrated sufficient effects of dECM at concentrations as little as 0.1-1.5 % w/v.^[45,47–49]^ Here it should be noted that a remaining limitation of dECM is obtaining and handling high concentrations, which can interfere with adequate digestion and pH adjustment.^[49]^ While this can be subject to optimization, increasing dECM concentrations will inevitably increase opaqueness, which in turn can interfere with imaging.

Our findings of increased levels of DNA damage with increasing stiffness across multiple cell types, support a role of ECM mechanics feeding into genomic instability. In accordance with this, our group has previously reported enrichment of DNA damage response pathways in MDA-MB-231 cells in 10 kPa alginate compared to 1 kPa or matrigel.^[63]^ We were able to further substantiate this using patient-derived TNBC cells, by showing that perturbation of FAK, a key mechanotransduction pathway, reverted the observed phenotype in stiff matrices. In line with this, a meta-analysis of genomic data across cancers revealed that cancers which arise in stiffer tissues display a higher mutational burden.^[28]^ Emerging literature further highlights the multiple, not mutually exclusive, pathways how confinement can lead to various forms of genomic instability. While some groups demonstrated how direct mechanical stress on the nucleus can lead to chromosome mis-segregation and replication stress, others report perinuclear mitochondrial localization and metabolic shifts leading to ATP, ROS or aldehyde accumulation which can change chromatin accessibility and induce DNA damage. ^[76–79]^

Our observations of lower lamin A/C levels in MDA cells and patient-derived TNBC in stiff matrices, align with the mesenchymal and metastatic features of TNBC cells. Indeed, previous studies have demonstrated that low lamin A/C levels, conferring more deformable nuclei, are associated with higher metastatic capacity and favor migration through confined spaces.^[26,80,81][18]^ The upregulation of lamin A/C in MCF7 on the other hand is in line with previous work, showing significant upregulation of lamin A/C in MCF7 encapsulated in 13 kPa compared to 2 kPa. ^[73]^ ^[81]^ Importantly studies in TNBC cell lines have indicated that various forms of nuclear compression, lead to replication stress-induced DNA damage in the absence of nuclear envelope rupture. Fibrosarcoma and fibroblasts on the other hand experienced nuclear rupture associated DNA damage. ^[82]^ In essence, this reinforces that cells subjected to the same stressors, can display distinct responses based on intrinsic (nuclear) properties.

Conducting additional exploratory analysis of BC patient cohorts, further substantiated this relationship between mechanical stress and DNA damage, revealing positive correlations between multiple mechanotransduction-related genes and DNA damage response-related genes (Supplementary Figure S7A). Ataxia telangiectasia and Rad3 related protein (ATR) constitutes a particularly compelling potential target as it is a central regulator in the DNA damage response, specifically in replication stress, and has also emerged to be a mechanosensitive protein. ^[83–85]^ Given that confinement can induce replication stress associated DNA damage in TNBC cells, implies a central role for ATR in maintaining genome integrity under mechanical stress.^[82]^ Interestingly, further analysis of BC cohorts demonstrated that both, FAK and ATR, tend to display elevated, albeit not significant, expression in TNBC (Supplementary Figure S7B). Moreover, TNBC patients expressing high levels of either mechanotransduction signature or DNA damage response signature experience worse distant metastasis free survival (Supplementary Figure S7C).

Based on our public database analysis, we experimentally investigated the potential of ATR inhibition (Ceralasertib) alone or in combination with the chemotherapeutic agent Gemcitabine, to sensitize BC cells experiencing mechanical stress-induced DNA damage. However, while the patient-derived TNBC cells displayed sensitivity to both reagents in Matrigel, no prominent effects were observed in Alg-dECM based matrices (Supplementary Figure S8A-B). This may be a result of distinct drug diffusion kinetics, with mesh size being a major determinant in this context. High molecular alginate networks may impose greater diffusional resistance than Matrigel, which is even more pronounced in stiff Alg-dECM at low N:T ratios. ^[86,87]^ In addition, Matrigel provides a highly growth promoting environment, making encapsulated cells potentially more susceptible to cytostatic drugs. Hence, the apparent lower sensitivity may reflect both the reduced proliferation activity and reduced effective drug exposure due to matrix mediated transport limitations. Overall, this data calls for further dose optimization and may require exploring targeting alternative pathways identified in our public data analysis (i.e. ATM, WEE1) (Supplementary Figure S7A). At this note, it should be emphasized that a remaining limitation of this platform is the incompatibility with high throughput screening. The rapid crosslinking kinetics of the material substantially impedes manual fabrication of large volumes and potentially require support with robotic platforms to facilitate this.

Finally, the compatibility with patient-derived cells demonstrated the clinical relevance of our model, which has multiple implications for future work. Two primary pillars of improving patient outcome are i) early detection and ii) personalized therapies. Tissue mechanics in combination with AI-algorithm-assisted pattern recognition are emerging as tangible, non-invasive biomarkers, that have been harnessed across distinct cancer entities.^[88–90]^ Our work suggests that assessment of BC patient’s lung tissue characteristics harbors underappreciated value as an additional readout to monitor disease course and guide therapeutic strategies. Next to enabling earlier detection, these readouts could also feed into optimized patient-based models. In this context, the multiple levels of tunability of our platform allow for the development of highly personalized models, incorporating not only patient cells but also the corresponding extracellular niche specific characteristics.

## 4. Conclusion

A tunable 3D bioengineered model recapitulating the biochemical and mechanical properties of healthy and metastatic lung niches revealed that increased matrix stiffness promotes ECM fiber organization, DNA damage and altered Lamin A/C levels in breast cancer cell lines and patient-derived TNBC cells. Pharmacological inhibition of FAK-mediated mechanotransduction reversed these effects, highlighting a mechanistic link between tissue mechanics and cancer cell phenotype. These findings reveal how altered tissue mechanics may contribute to breast cancer progression and provide a platform for patient-derived models.

## 5. Experimental Section

### Norbornene & tetrazine modified alginate

HMW (MW> 200 kDa, MVG Pronova, 42000101) or LMW (MW> 75 kDa, VLVG Pronova, 4200501), high guluronic acid sodium alginate was coupled to 5-norbornene-2-methylamine (N, TCl Chemicals, N0907) and tetrazine (T, Conju-probe, CP-6021), with a theoretical degree of substitution (DS_theo_) of 500 and 300 molecules per alginate chain, respectively. Upon dissolving alginate at 1 % w/v in 0.1 M 2-(N-morpholino)ethanesulfonic acid (MES; Sigma Aldrich, M5287), 0.3 M sodium chloride (NaCl; BioChemica, 131659.1211) at a pH of 6.5, N-hydroxysuccinimide (NHS, Sigma-Aldrich,130672) and 1-ethyl-3-(3-dimethylaminopropyl)-carbodiimide hydrochloride (EDC, Sigma-Aldrich, E7750) were added drop-wise while stirring. Then, either N or T was added, yielding a final reaction concentration of 0.6 % w/v. After 20 h, the reaction was quenched by adding hydroxylamine hydrochlorine (HCl, Sigma-Aldrich, 26103) for 30 min followed by 30 min centrifugation at 10k rpm to remove residual/unreacted N or T. The solutions were then dialyzed against a descending NaCl gradient with 10x phosphate buffered saline (PBS, Gibco, 14200-0670) and deionized (DI) water using dialysis membranes (Spectra/Por 6 molecular cut-off 3.5kD, 15370732) with dialysis changes every 3-4 hours prior to overnight dialysis. This was continued for 3 days. After the final dialysis, activated charcoal (Sigma-Aldrich, C9157) filtration was performed prior to sterile filtering (0.22 μm; Steriflip-GP; Merck, SCGP00525) and overnight storage at −20°C. The following day, samples were lyophilized for 3 days and stored at −20°C until usage.

### Nuclear magnetic resonance

The degree of substitution (DS_actual_) of N-Alg and T-Alg was determined by performing nuclear magnetic resonance (NMR; 64 scans, NMR – Bruker Ultra Shield Plus 500 MHz) of 1.5 % w/v N- and T-alginate solutions in deuterium oxide (D2O, Millipore, 1.03428.0009). Resulting NMR spectra were analyzed using MestreNova software (version 11.02) to calculate each DS_actual_. In brief, upon averaging the N peak area at 6.2-5.8 ppm and T peak area at 10.2 ppm, the resulting values were divided by the area of the Alg peak (G-1) between 5.2-4.8 ppm.

### Decellularization of porcine lung tissue

Porcine lung tissue was obtained from the local animal facility (Biogipuzkoa), dissected and stored at −80 °C until decellularization. Upon thawing, the tissue was minced into small pieces (1 mm^3^) using surgical scalpels. The tissue was then transferred into 50 mL falcons, until the cone mark of the tube to ensure sufficient exposure of the tissue to the reagents. Unless stated otherwise, the reagents were filled until the 45 mL mark and the following steps were performed on a rotator. First the tissue was subjected to 0.1 % Triton X-100 (Sigma-Aldrich, T9284) for 30 minutes (min), followed by 30 min 1x PBS incubation and a subsequent overnight treatment with 2 % w/v sodium deoxycholate (Sigma-Aldrich, D6750) at 4 °C. The following day, washes with 1x PBS in 15 min intervals for 2 h were performed, which was then repeated with DI water.

The tissue was then treated with 1 M NaCl (Fisher Scientific, 10428420) for 30 min, washed with DI water for 30 min, 1 % Triton for 30 min and finally 1x PBS overnight at room temperature. The next day the tissue was subjected to deoxyribonuclease treatment (30 units/mL, Bovine DNAse I, Sigma-Aldrich, D5319) in 30 mL 1.5 mM magnesium chloride (Cl_2_, Sigma-Aldrich, M8266) at 37 °C overnight. On the final day of decellularization, the tissue was sterilized with 0.1 % v/v peracetic acid (PA, Sigma-Aldrich, 1072221000) and 4 % v/v ethanol (Scharlau, ET0002005P) for 2 h, washed twice with 1x PBS for 15 min, followed by 2 additional washes with DI water for 15 min prior to overnight storage at −80 °C. The following day, the decellularized tissue was lyophilized for 3 days and ground into powder. For pre-gel dECM stock solutions, powder was digested at RT with pepsin (0.2 mg/mg dECM, Sigma-Aldrich, P7012-1G) in 0.5 M acetic acid (Sigma-Aldrich, 695092) for 48 h. Digestion was terminated by adjusting the pH to 7.25-7.45 on ice, using 2-5 M sodium hydroxide (NaOH, Sigma-Aldrich, 221465). Pre-gel stock solutions were stored at 4 °C. While for characterization purposes, three independent samples derived from three donors were used, a single donor was used for the herein performed cell-based experiments.

### Histological assessment of native lung and dECM

Paraffin (Sigma-Aldrich, MKCT5465) was melted the day prior to embedding at 80°C. The following day, fixed native and decellularized tissue samples were dehydrated in ascending ethanol series (2 x 50 % v/v, 2 x 70 % v/v, 3 x 96 % v/v) for 20 min each step, followed by 2 x 30 min xylene (Bio Optica, 06-1304Q) incubations. Samples were then transferred to a pre-warmed xylene-paraffin (1:1) for 30 min and subsequently embedded in paraffin 2x 30 min. Tissue blocks were allowed to solidify overnight and cut into 5 µm sections using a Microtome (Leica HM 355S). Sections were deparaffinized and rehydrated in a descending ethanol series, permeabilized and stained with Hematoxylin (Bio Optica, 05-06004/L) and Eosin (Sigma-Aldrich, HT11016) (H&E), Picrosirius red, Venhoerf Van Giesson and DAPI (ThermoFisher, 62248) as described previously.^[91]^ Finally, sections were mounted in DPX mounting medium (Fisher Scientific, 10383191) and imaged using an inverted epifluorescence microscope AxioObserver 7 (Zeiss) microscope equipped with a polarizer and a EC Plan-Neofluar 10x/0.30 NA Ph1 Objective.

### DNA quantification of native lung and dECM

DNA content of native and decellularized lung tissue was quantified using Quanti-iT^TM^ PicoGreen^TM^ kits (Invitrogen, P7589). In brief, tissue samples were digested for 72 h at 37 °C in 500 µL digestion buffer containing 6 mM EDTA (Invitrogen, AM9260), 6 mM L-Cysteine (Sigma-Aldrich, 168149) and 4 units/mL Papain (Sigma-Aldrich, P4762). Afterwards, samples were incubated for 1 h at 65 °C, subsequently centrifuged at 10k rpm and supernatants were collected. DNA was precipitated for 1 h on ice. Samples were centrifuged for 30 min at 0°C at 10k rpm and supernatants were discarded. Pellets were resuspended in 70 % ethanol and centrifuged at maximum speed for 2 min, this was repeated 3x. Pellets were left to dry before resuspension in 1x TE buffer to prepare serial dilutions of both tissue samples and a DNA standard to establish standard curves. Upon adding 1:200 diluted fluorescent reagent provided by the quantification kit, absorbance was measured, 492 nm excitation and 535 nm emission.

### Proteomic analysis of native lung and dECM

To analyze protein composition of native porcine lung tissue and corresponding pre-gel lung dECM (that is the decellularized, lyophilized and pepsin-digested stock solution), label-free liquid chromatography-mass spectrometry (LC-MS/MS) was conducted at the proteomic platform at CIC bioGUNE (Bilbao, Spain). In brief, protein samples of three independent biological replicates were extracted in a buffer containing 7 M urea, 2 M thiourea, 4 % w/v 3-[(3-cholamidopropyl) dimethylammonio]-1-propanesulfonate (CHAPS) and 5 mM dithiotreitol (DTT). Subsequently samples were digested following the filter-aided FASP protocol described by Wisniewski et al. with minor modifications, as indicated in the following.^[92]^ Trypsin was added to a trypsin:protein ratio of 1:20, and the mixture was incubated overnight at 37 °C, dried out in a rotation vacuum concentrator (RVC2 25 speedvac, Christ), and resuspended in 0.1 % formic acid (FA).

Samples were analyzed in a hybrid trapped ion mobility spectrometry – quadrupole time of flight mass spectrometer (timsTOF HT, Bruker Daltonics) coupled online to a an Evosep ONE liquid chromatograph (EVOSEP). Purified phosphopeptides (20 % w/v of the purified material) were directly loaded onto the Evosep ONE and resolved using the 30 samples-per-day protocol. Protein identification and quantification was carried out using Fragpipe software (https://fragpipe.nesvilab.org/).^[93]^ Searches were carried out against a database consisting of Sus scrofa (porcine) entries (Uniprot/Swissprot), with precursor and fragment tolerances of 20 ppm and 0.05 Da. Carbamidomethylation of cysteines was considered as fixed modification, and oxidation of methionine was considered as variable. A false discovery rate (FDR) <1 % at peptide level was applied as a significance cutoff.

Protein abundances obtained from Fragpipe (MaxLFQ Intensities) were loaded onto Perseus platform (v.2.1.2).^[94]^ These values were log2 transformed and only those proteins present in at least 70 % of the samples of any of the groups under analysis were considered for further analyses. Missing values were imputed using the QRILC method available in the Perseus version (https://www.maxquant.org/perseus/) used under default parameters.

Publicly available healthy total human lung (https://www.ebi.ac.uk/pride/archive/projects/PXD010154) and compared to porcine native and dECM lung proteome via a web-based Venn Diagram tool (Evenn).^[56,95]^ Gene ontology (STRING) analysis was performed to determine molecular functions of terms enriched in lung dECM with and FDR of 0.5 set as the level of significance.^[96]^ In addition, enriched reactome pathways were determined and compared to publicly available human lung ECM and matrigel proteomic data sets applying an FDR of 0.5 as the level of significance.^[45,58,96]^ Moreover, we mapped these genes to the Matrisome database, a human based ECM catalogue, to further annotate ECM specific proteins.^[97]^

### Click alginate-dECM hydrogel casting

Modified N-Alg and T-Alg was weighed and dissolved separately in 1x PBS overnight at RT on a shaker at 500 rpm. The following day formulations were combined using positive displacement pipettes (Gilson) to ensure accuracy with viscous materials. First lung dECM was added to the Alg component with the lower volume (T-Alg in 4.0 N:T ratios, and N-Alg in 0.5 N:T ratios) at a final concentration of 0.5 % w/v dECM and mixed thoroughly to ensure a homogenous distribution. Afterwards, the remaining Alg component with the higher volume (N-Alg in 4.0 N:T and T-Alg in 0.5 N:T formulations) was added with the total final concentration of Alg being 2.5 % w/v. Covalent click crosslinking via inverse electron demand Diels Alder reaction of N-T Alg takes place rapidly, thus mixing via pipetting was performed at a pace that allowed homogenous distribution of each component prior to crosslinking while limiting introducing air bubbles. If required, centrifugation was performed between mixing steps to remove air-bubbles. The final formulation was then transferred to house made silicone molds of 1 mm height and 4 mm diameter, sandwiched between sigma coated (Sigma-Aldrich, SL2) glasses to yield even hydrogel surfaces and prevent adhesion of the crosslinked hydrogels to the glasses. The molds were transferred to a cell incubator at 37 °C for 30 min to allow physical collagen crosslinks to form within dECM. Finally, crosslinked gels were transferred to 96 well plates with 1x PBS and allowed to equilibrate prior to mechanical characterization.

### Rheology

Storage (G’), loss (G’’) and complex (G*) modulus of hydrogels were determined with a rheometer (Anton Paar MCR302) equipped with a parallel sandblasted plate geometry of 8 mm (Anton Paar, PP08). N-T-Alg hydrogels with and without dECM at various concentrations, MWs and Alg:dECM as well as N:T ratios were casted as described above using 1 mm thick and 8 mm diameter custom made silicon molds. Gels were allowed to equilibrate overnight in 1x PBS. Upon initial contact with the gel surface, a compression of 10 % at the corresponding height of the hydrogel was applied. Subsequently, frequency sweeps were performed from 0.03 to 30 Hz at 0.3 % shear strain at RT. To derive the elastic moduli (E) the following formula was applied: E = 2G*(1 + ϑ) where G* falls within the linear viscoelastic region and where ϑ is he Poisson’s ratio with a value of 0.5 for hydrogels.^[98]^

### Confocal reflectance imaging and analysis

Collagen fiber morphology and architecture in lung dECM containing hydrogels was determined using a confocal microscope (LSM880, Zeiss) in reflectance mode. Gels were imaged with a C-Apochromat 40x/1.2 NA Korr FCS M27 objective, a 488 nm argon laser at 0,1 % and PMT detector at 488 nm excitation and 490 nm emission, to obtain Z-stacks of 10 µm at 1 µm intervals.

To determine collagen fiber morphology, Z-stacks were projected and analyzed using CT-FIRE analyses (version 3.0 Beta for windows, 2020).^[52]^ Fiber coherency and orientation maps were obtained using OrientationJ Plugin for ImageJ (version 1.54).^[99]^

### Patient sample collection

Samples and data from patients included in this study were provided by the Basque Biobank www.biobancovasco.bioef.eus (National Registry of Biobanks (B.0000140), integrated in the Platform ISCIII Biomodels and Biobanks (PT23/00013). Sample management, clinical data and all procedures were performed under the ethical approval from the Ethics Committee of the Basque Country (CEIm-E, Code PI+CES-BIOEF 2024-13).

### Paraffin-embedded BC lung metastatic tissue sections

Formalin-fixed paraffin-embedded (FFPE) tissue blocks were processed and prepared by the corresponding Pathology Departments and subsequently provided by the corresponding Basque Biobank nodes from HU Basurto and HU Cruces. Tissue fixation, paraffin embedding, and sectioning were performed according to the standard operating procedures of the Pathology Departments and the Basque Biobank. Briefly, biopsy samples were sectioned, and the most representative ones were selected, taking tumor margins into account. Samples were fixed in 10% formalin, dehydrated in 50% alcohol, followed by xylene washes, prior to paraffin embedding. Resulting paraffin blocks were then cut in 3 µm thick sections. H&E and immunohistochemistry (IHC) stained slides by the Pathology Department were digitalized in our lab and returned to their archives.

### Multiphoton imaging and analysis

SHG and TPEF images of unstained slides of paraffin embedded tissue samples were acquired using a laser scanning confocal microscope (LSM880, Zeiss). Samples were excited using a Mai-Tai DeepSee laser set to a wavelength of 800 nm wavelength and at 1.8 % power. A Plan-Apochromat 20x/0.8 NA M27 objective at zoom 1x was used to acquire overview tiles and a zoom of 3x was applied to acquire images for analysis within the tumor region. Detection wavelength of SHG and TPEF via non-descanned detectors were set at 390-415 nm and 465-515 nm, respectively, using 2 non-descanned GaAsP detectors. To analyze fiber architecture, both channels were subjected to CTfire analysis (version 3.0 Beta for windows, 2020).^[52]^

### BC cell line culture

Wild type (WT) and previously established genetically modified FUCCI2 MDA-MB-231 (HTB-26; American Type Culture Collection (ATCC), Lot. 62657852) and MCF7 (HTB-22; ATCC, Lot. 63288596) were cultured in low-glucose Dulbeccos modified Eagles medium (DMEM; Sigma-Aldrich, D6046) with 10 % v/v fetal bovine serum (FBS, Gibco A5256701) 1 % v/v penicillin/streptomycin (Teknovas, A315140122) with additional 1 % v/v Glutamax (Gibco, 25030024) to MCF7 media.^[62–64]^ Cells were passaged twice a week and maintained at 37 °C 5 % CO_2_. Cells were tested for mycoplasma by PCR-based mycoplasma detection assay every month and consistently confirmed negative.

### Patient-derived triple negative breast cancer cell culture

Patient-derived biopsies were obtained in the diagnostic procedure according to clinical standards of the Breast Radiology Service from Hospital Donostia and under ethical approval from the Ethics Committee of the Basque Country (CEIm-E, Code PI+CES-BIOEF 2024-13). Extra core needle biopsies (CNB) were extracted for this project and recovered into a centrifuge tube in MACS Tissue Storage Solution (MACS Milteny, 130-100-008). For this study we opted for a CNB of a TNBC patient, identified based on information by trained pathologists. The CNB was preserved at 4°C and processed within 3 hours after extraction. The CNB was placed in a sterile 10 cm-dish. Fatty as well as visually recognizable non-tumoral tissue was removed by using sterile disposable blades (Swann_morton, ABH5011). The remaining tumor tissue was further minced to approximately 1 mm^3^ fragments. A volume of 100 μL of 10x Collagenase/Hyaluronidase (STEMCELL technologies, 07912) and 900 μL of Basic Media consisting of Advanced DMEM-F12 (Gibco, 12634010) with HEPES 1 % v/v (Gibco, 15630080), Penicillin-Streptomycin 1 % v/v (ThermoFisher, 15140122) and L-Glutamine (Gibco, 25030024) was added to the fragments, followed by an incubation at 37°C and 800 rpm in a thermo-shaker for 45 minutes to one hour. Afterwards, the sample was filtered through a pre-wetted 100 μm cell strainer (Corning, CLS431752-50EA) into a new 50 mL centrifuge tube. Remnant tissue was pipetted up and down in order to further dissociate it and filtered again through the strainer. The filtered material was then centrifuged at 500 g for 5 min at 8 °C and, followed by a red blood cell lysis with ACK Lysing Buffer (Gibco,A1049201) for 3 min in a roller at RT. Cells were washed with 1x DPBS and centrifuged again prior to resuspending the pellet in Complete Medium (Basic Medium + 10 ng/mL EGF (PeproTech, AF-100-15-100UG) + 5 µM Y-27632 (MedChemExpress, HY-10583), 1 µg/mL Hydrocortisone (Sigma-Aldrich, H088-1G) + 100 µg/mL Primocin (InvivoGen, ant-pm-05) + 10 % v/v FBS (Gibco, A5256701) and being seeded in one well in a 6-well plate in adherent conditions. For the first two weeks to 4 weeks, FBS was kept in the medium in order to help initial proliferation but then omitted to prevent fibroblasts overgrowth. Medium was changed two to three times a week and, under subconfluent conditions, culture was passaged into larger surfaces. Medium formulations and procedure were adapted from Bock and colleagues.^[100]^ When the culture got to three 75 cm^2^ flasks, it was finally frozen down in Cryostor CS-10 freezing medium (STEMCELL Technologies, 07930). When needed for experiments, cells were thawed and kept in the same described conditions until needed for hydrogel encapsulation upon counting. The patient-derived culture was routinely tested for mycoplasma contamination and was confirmed to be negative in all experiments included in this study.

### Cell encapsulation

For cell encapsulation, hydrogels were prepared as described above. Cells were detached using 0.5 % trypsin EDTA and counted with an automated cell counter (Countess 3, Invitrogen). Upon centrifugation at 500 g for 5 min, cells were resuspended to yield single cell suspensions at a density of 5 x 10^5^ cells/mL in the final hydrogel formulation. Then the required cell suspension volume was added to the Alg component with the lower volume (and gently but thoroughly combined with the cell suspension. Afterwards either lung dECM or 1x PBS was added before introducing the remaining Alg component. Crosslinked gels were then transferred to 96 well plates with media and maintained in a cell incubator. Media was changed every 2-3 days.

### Cell viability

To assess viability, live dead assays using calcein AM (AATbioquest, 2202) and ethidium homodimer-1 (EthD-1, Sigma-Aldrich, E1903) were performed on day 1, 7 and 14. Gels were washed with 1x PBS and incubated at 37 °C for 10 min with 4 mM calcein AM and 2 mM Ethd-1. Gels (n=3 per condition) were washed with 1xPBS and placed on a microscope slide for imaging using an AxioObserver 7 (Zeiss). Z-stacks of 200 µm were taken in the center of the gel in 10 µm Z-steps with an EC Plan-Neofluar 10x/0.30 NA Ph1 objective. Z-stacks were analyzed to quantify viability via custom-trained deep learning models using Dragonfly imaging software (v.2.2, Object Research Systems (ORS) Inc. Montreal, Canada 20218).

### FUCCI imaging & analysis

Longitudinal imaging of FUCCI modified MCF7 and MDA cells at day 1, 7 and 14 was performed using a AxioObserver 7 (Zeiss). Z-stacks in the center of the gels (n=3 per condition) spanning 200 µm with 10 µm Z-steps were acquired using an EC Plan-Neofluar 10x/ 0.30 NA Ph1objective. For 3D FUCCI analysis, mCherry and mVenus signal was segmented and quantified using Dragonfly software (version 2.2, Object Research Systems (ORS) Inc.,Montreal, Canada, 20218) via custom-trained deep learning models.

### DAPI-phalloidin immunofluorescence imaging & analysis

Gels (n=3 per condition) were washed in 1xPBS for 5 min, fixed in 4 % v/v paraformaldehyde for 15 min, washed in 3 % w/v BSA 2 x for 20 min and permeabilized with 0.03% v/v Triton X-100 for 30 min. Afterwards, gels were washed with 3 % w/v BSA 3 x 20 min, 3 % w/v BSA with and incubated with DAPI (1:1000; Merck, MBD0015) and phalloidin ATTO 550 (1:100; Sigma-Aldrich, 19083) in 3 % w/v BSA overnight. The following day gels were washed in 1x PBS 3x for 5 min. Z-stacks of 100 µm with 1 µm z-steps were taken using a AxioObserver 7 (Zeiss) equipped with an apotome and an EC Plan-Neofluar 10x/0.30 NA Ph1 objective. 3D cluster morphology was quantified using Dragonfly software (version 2.2, Object Research Systems (ORS) Inc.,Montreal, Canada, 20218) via custom-trained model using the guided workflow Segmentation Wizard tool. Multiple frames (3-4) were manually labeled to generate clean data via a Random-Forest quick start model at pre-defined default training settings. The final segmentation model was trained using U-Net machine learning classifier also at default settings (patch size 64; epoch number 100; stride ratio 0.25; batch size 256; loss function OrsDiceLoss; optimization algorithm Adelta). Upon cluster segmentation, clusters located at the boundary slices (beginning and end of z-stacks) or image border were manually excluded from analysis to prevent underestimation of cluster volumes due to truncation artifacts. A volumetric lower-bound cut-off of 500 µm^3^ (≈9.8 equivalent spherical diameter), which sits below the average size of human breast cancer cells, was applied to filter out units/artifacts below single cell level such as phalloidin debris, dead cells or sub-cellular fragments.

### Lamin A/C and yH2AX immunofluorescence imaging & analysis

Co-staining’s of DAPI-Phalloidin with lamin A/C and yH2AX were performed by washing gels in 1x PBS for 5 min, fixed in 4 % v/v paraformaldehyde for 15 min, washed in 3 % w/v BSA 2x for 20 min and permeabilized with 0.03 % v/v Triton X-100 for 30 min. Afterwards, gels were washed with 2 % w/v BSA 2 x 20 min, 2 % w/v BSA with 0.01 % v/v Tween-20 for 20 min and incubated overnight with primary antibodies: mouse-monoclonal lamin A/C (1:500; Cell Signaling Technologies, 4777S) and yH2AX rabbit-monoclonal antibody (1:500, Abcam, AB81299) in 2 % BSA over night at 4 °C. The following day, gels were washed with 3 % w/v BSA 2 x 20 minutes and stained with secondary donkey anti-mouse AF647 (1:500; Cell Signaling Technologies, 4410), donkey anti-rabbit AF488 (1:500, Cell Signaling Technologies, 4412) as well as DAPI (1:1000; Merck, MBD0015) and phalloidin ATTO 550 (1:100, Sigma-Aldrich, 19083) in 2 % w/v BSA overnight at 4°C. Finally, gels were washed in 1x PBS 3 x for 5 min. For quantification, Z-stacks of 80 µm with 2 µm Z-steps were acquired using an AxioObserver 7 (Zeiss) with an LD Plan-Neofluar 40x/0.6 NA Korr Ph2 M27 objective.

To obtain lamin A/C and yH2AX intensities, Z-stacks were projected and images were segmented for nuclei (Dapi) using ImageJ (version 1.54) via automatic thresholding. Dapi based nuclear ROIs were then semi-automatically defined using the magic wand tracing tool.^[99]^ Within these nuclear ROIs, corresponding intensity in lamin A/C and yH2AX channels were measured. Background corrections were applied by subtracting intensities measured within cell free ROIs in the respective channels.

### Analysis of mechanotransduction genes & DNA damage response genes

To assess and guide further experimental design, expression of key mechanotransduction and DNA damage response genes, as well as correlation analysis was conducted using GEPIA (http://gepia2.cancer-pku.cn).^[101]^ Finally, the prognostic impact of the signature genes was evaluated via survival analysis of TNBC cohort for distant metastasis free survival (DMFS), using the Kaplan-Meier plotter online tool (http://kmplotter.com).^[102]^

### FAK inhibition

Defactinib (VS-6063; PF-04554878 MedChemExpress, 1073154-85-4), a clinically relevant FAK-inhibitor was chosen to interfere with mechanotransduction. Working stocks of 1 mM, 2.5 mM and 5 mM, were prepared from a 10 mM master stock in DMSO to yield equivalent DMSO concentration across all tested concentrations. Upon encapsulating patient-derived triple negative BC cells in Alg:dECM at a density of 500.000 cells/mL, cells were treated at a concentration range of 1 µM, 2.5 µM and 5 µM or DMSO (0.1% v/v) as control. Fresh media with Defactinib was prepared prior to media change every 3 days for 14 days. At the endpoint of the culture, cells were fixed and stained with DAPI, phalloidin, yH2AX and Lamin A/C as described above.

### Drug testing

Ceralasertib (AZD6738, 1352226-88-0, MedChemExpress), a selective ATR-inhibitior, under active clinical investigation was chosen to assess whether elevated DNA damage could be therapeutically exploited in combination with Gemcitabine (LY 188011 hydrochloride, 122111-03-9, MedChemExpress). Initial dose determinations were carried out by exposing patient-derived triple negative BC cells (500.000 cells/mL) encapsulated in Matrigel (356237, Corning, ∼10 mg/ml) for 72 hours, to concentrations of 100 nM, 250 nM, 500 nM, 1 µM, 2.5 µM and 5 µM of Ceralasertib or 50 nM, 100 nM, 250 nM, 500 nM, 1 µM, 2.5 µM and 5 µM of Gemcitabine for subsequent 48 hours. Viability assays were performed as described above and informed the determination of appropriate concentrations for each compound. For the final experimental setup, patient-derived triple negative BC cells were encapsulated in stiff and soft Alg-dECM for 5 days, and exposed to 5 µM Ceralasertib and 5 µM Gemcitabine individually and in combination for subsequent 3 days.

### Statistical analysis

Bar graphs display mean values and standard deviation, while dot graphs display the median with interquartile range. All statistical analysis was carried out using GraphPad Prism (version 10.5.0, GraphPad Software Inc., Boston, Massachusetts USA). Data was assessed for normality based on Shapiro Wilk test and equal variances were determined based on F-tests. To compare two groups, two tailed Student’s t-test or Mann-Whitney U-test were used to compare groups with normal or not-normal distribution, respectively.

## Acknowledgements

The authors are grateful for the support of Irantzu Llarena at CIC biomaGUNE with advanced optical imaging and the Proteomics platform at CIC bioGUNE for the guidance during proteomic analysis. We would also like to express our gratitude to Unai Heras for explaining rheological concepts and instructions on 3D image analysis. We are grateful to the MATRIXinCANCER clinical network, with special thanks to Amaia de la Maza and Leire Romarate as Clinical Research Coordinators, and data managers from Biogipuzkoa and Biobizkaia. We want to particularly acknowledge the patients and the Basque Biobank www.biobancovasco.bioef.eus integrated in the Platform ISCIII Biomodels and Biobanks (PT23/00013) for their collaboration. We further acknowledge the Breast Radiology Department members from HU Donostia; and the Pathology Department members from HU Donostia, HU Basurto and HU Cruces, especially their respective heads: Irune Ruiz, Goikoane Cancho and Leire Andrés.

L. Fallert acknowledges Investigo programme, funded by European Union–NextGenerationEU as part of Spain’s Recovery, Transformation and Resilience Plan as well as the Collaborative PhD Fellowship CICbiomaGUNE Program 2022 (CICBMG_PhD_03_2022). A. Cipitria would like to acknowledge funding from Fundación Científica Asociación Española Contra el Cáncer (grant LABAE223466CIPI), from the Spanish Ministry of Science and Innovation (MCIN/AEI/10.13039/501100011033/FEDER UE, through grants #PID2021-123013OB-I00 and #PID2025-169199OB-I00), from Programa FORTALECE - IIS Biodonostia (FORT23/00026) and from the European Research Council Consolidator Grant (DORMATRIX, 101123883). D. Jimenez de Aberasturi would like to acknowledge funding from the Spanish Ministry of Science and Innovation (MCIN/AEI/10.13039/501100011033; “ERDF A way of making Europe and European Union NextGenerationEU/PRTR”, through grants #PID2022-143248OB-I00 and #CNS2022-135941), from the Basque Government, R&D Projects in Health 2025 (Grant no. 2025333046). She also thanks the Spanish Government for “Ramon y Cajal” Fellowship (#RYC2022-037249-I). A. Cipitria and D. Jimenez de Aberasturi also acknowledge funding from IKERBASQUE Basque Foundation for Science.

## Data Availability Statement

All data supporting this article including raw proteomic data, microscope images of all replicates and Dragonfly models are available at Zenodo repository (10.5281/zenodo.2253065).

Received: ((will be filled in by the editorial staff))

Revised: ((will be filled in by the editorial staff))

Published online: ((will be filled in by the editorial staff))

