## Supplementary Information for "Lung-mimicking click alginate-dECM model of breast cancer lung metastasis reveals the role of ECM mechanics in tumor growth dynamics and genomic instability"

- Supplementary Figure S1: BC patient lung metastasis samples.
- Supplementary Table S1: Clinicopathological BC patient data and sample information.
- Supplementary Figure S2: Comparative proteomic analysis.
- Supplementary Figure S3: Low molecular weight alginate characterization.
- Supplementary Figure S4: High molecular weight alginate characterization.
- Supplementary Figure S5: Biocompatibility of Alg and Alg-dECM.
- Supplementary Figure S6: Nuclear and genomic integrity in stiff and soft alginate.
- Supplementary Figure S7: Analysis of mechanotransduction and DNA damage in BC patients.
- Supplementary Figure S8: Drug testing with patient-derived TNBC cells.

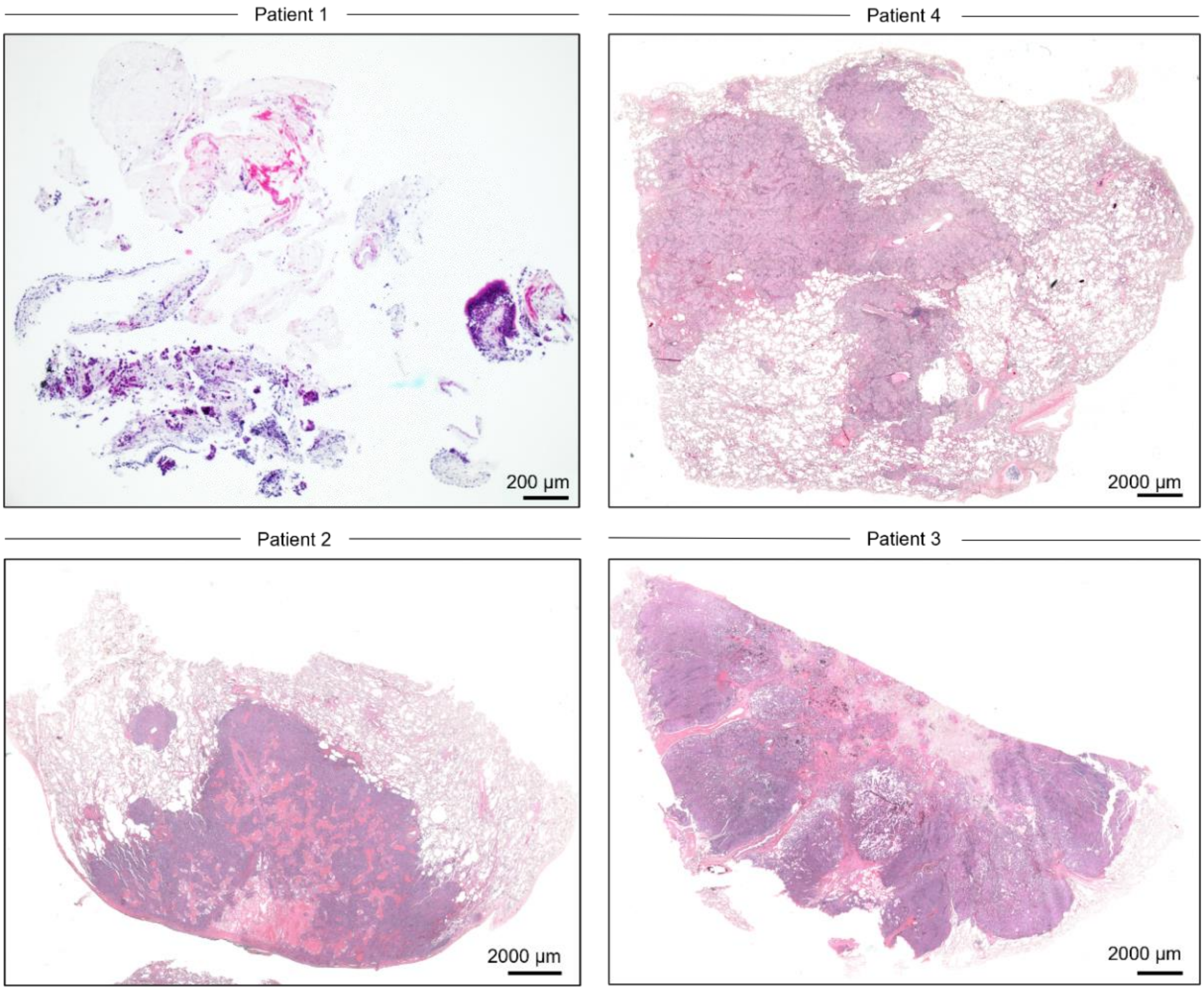


**Supplementary Figure S1.** BC patient lung metastasis samples. A) Bright field images of H&E stained FFPE sections.

**Supplementary Table S1.** Clinicopathological BC patient data and sample information

| | Patient 1 | Patient 2 | Patient 3 | Patient 4 | Patient 5 |
|---|---|---|---|---|---|
| **Sample type** | FFPE | FFPE | FFPE | FFPE | Fresh biopsy (cells) |
| **Age (at time of diagnosis)** | 56 | 52 | 53 | 37 | 71 |
| **Sex** | Female | Female | Female | Female | Female |
| **Histological subtype** | TNBC | HER2+ | ER+/Luminal-like | HER2+ | TNBC |
| **Sample origin** | Lung | Lung | Lung | Lung | Breast |
| **Clinical stage at diagnosis** | Not specified | Not specified | 0 | Not specified | III (localized) |
| **Menopausal status (at time of diagnosis)** | Not specified | Postmenopausal | Postmenopausal | Premenopausal | Postmenopausal |
| **ER (%)** | 0 | 0 | 100 | 0 | 0 |
| **PR (%)** | 0 | 0 | 40 | 0 | 0 |
| **HER2 IHC Score** | 1 | 3 | 1 | 3 | 1 |
| **Histological grade** | Not specified | 2 | Not specified | Not specified | 3 |
| **Ki67 (%)** | Not specified | 25 | 12 | 36 | 70 |

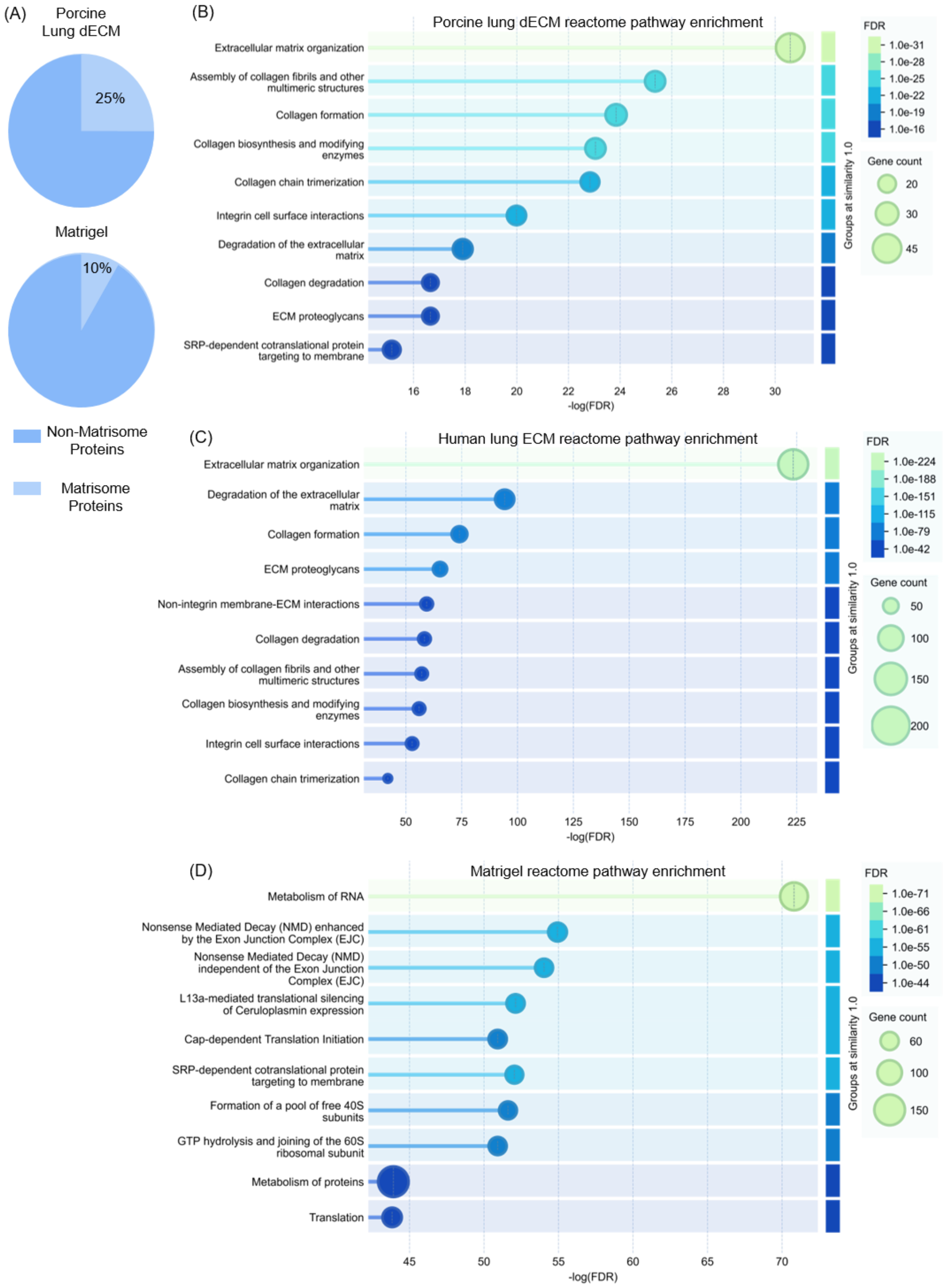


**Supplementary Figure S2.** Comparative proteomic analysis. A) Pie charts displaying percentage of matrisome and non-matrisome related proteins of porcine lung dECM and Matrigel. B) Reactome pathway analysis of porcine lung dECM. C) Human lung ECM and D)

Matrigel showing the top 10 most significantly enriched reactome pathways based on STRING protein interaction analysis. Pathways were filtered based on a false discovery rate (FDR) of >0.05 and the graphs show log-transformed FDR values, with node size representing the number of genes associated with each Gene Ontology term, and node color intensity reflecting the -log(FDR).

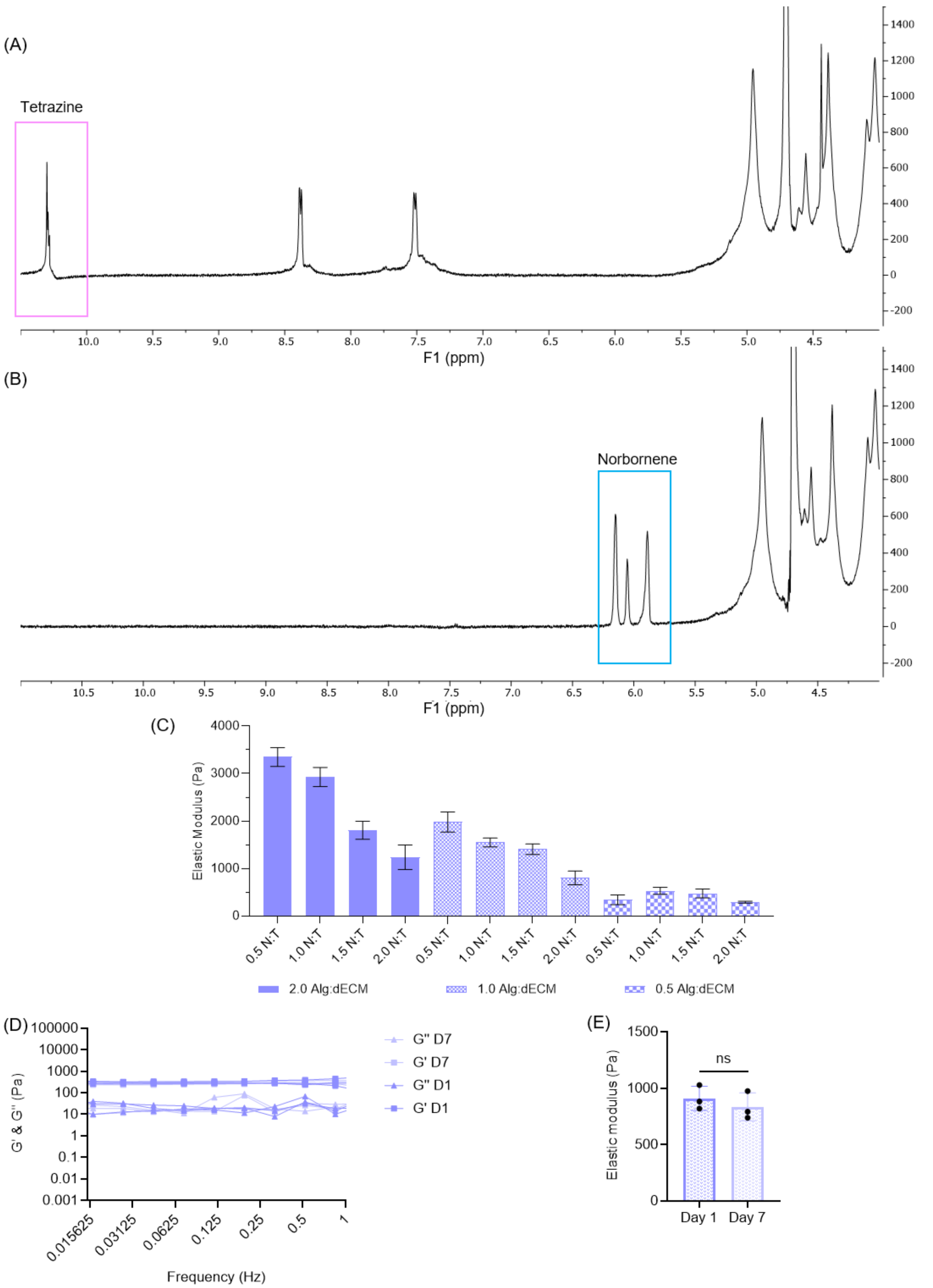


**Supplementary Figure S3**. Low molecular weight alginate characterization. A) NMR spectra of tetrazine (peak of interest for DS calculation at 10.2 ppm) and B) norbornene (peaks at 6.2-5.8 ppm) modified low molecular weight alginate (75 kDa) with a DS real of 34.4 and 31.7

respectively. C) Mechanical properties of Alg-dECM at a constant total polymer concentration of 2% w/v with varying N:T ratios and Alg:dECM ratios. D) Frequency sweep and E) corresponding elastic moduli of 2% w/v Alg with 2% w/v dECM, at N:T ratio of 1, with cells, at day 1 and day 7. n = 3 gels per conditions. Bar graphs display means with standard deviation. Mann-Whitney U test.

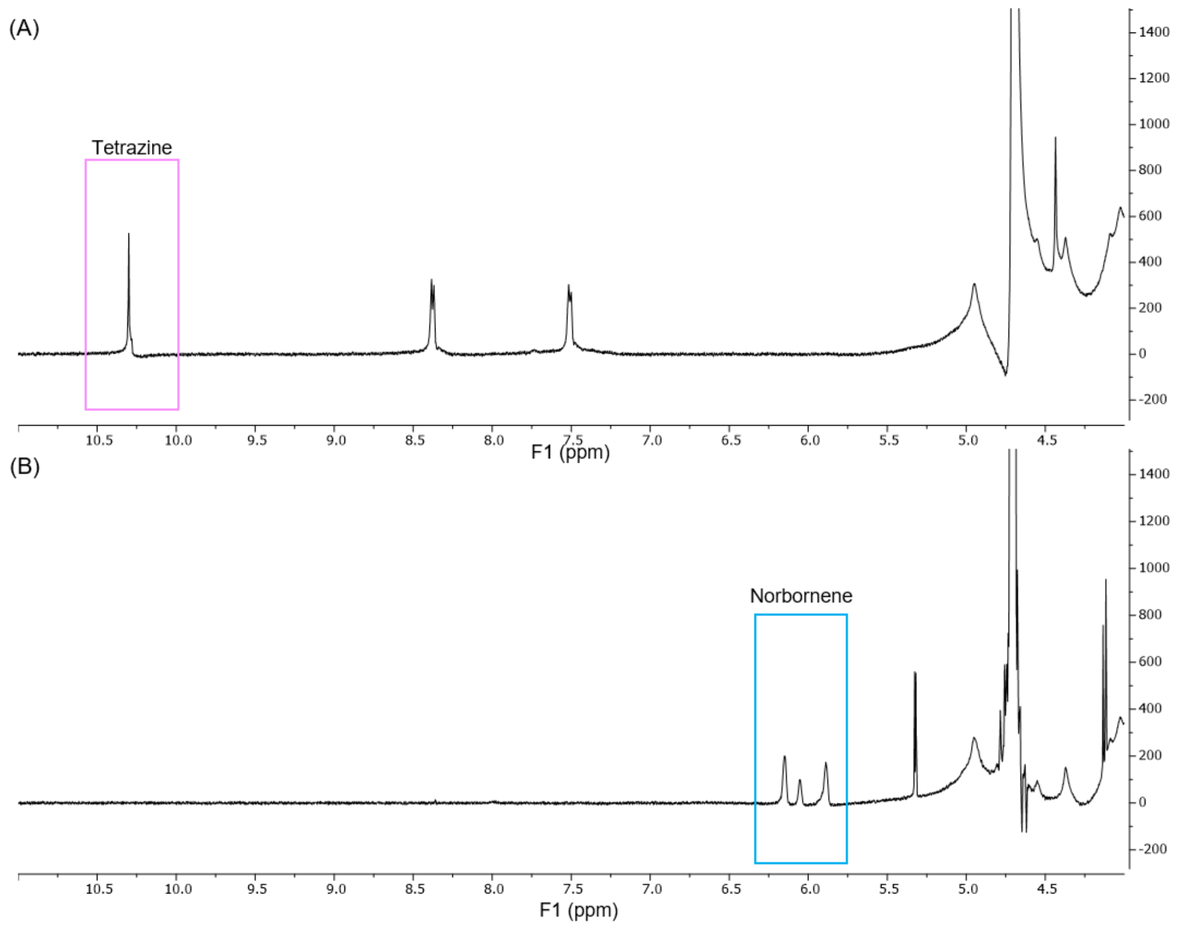


**Supplementary Figure S4.** High molecular weight alginate characterization. A) NMR spectra of modified high molecular weight alginate (200 kDa) modified with tetrazine (peak of interest for DS calculation at 10.2 ppm) and B) norbornene (peaks at 6.2-5.8pppm). DS real of 121.2 and 99.3 respectively.

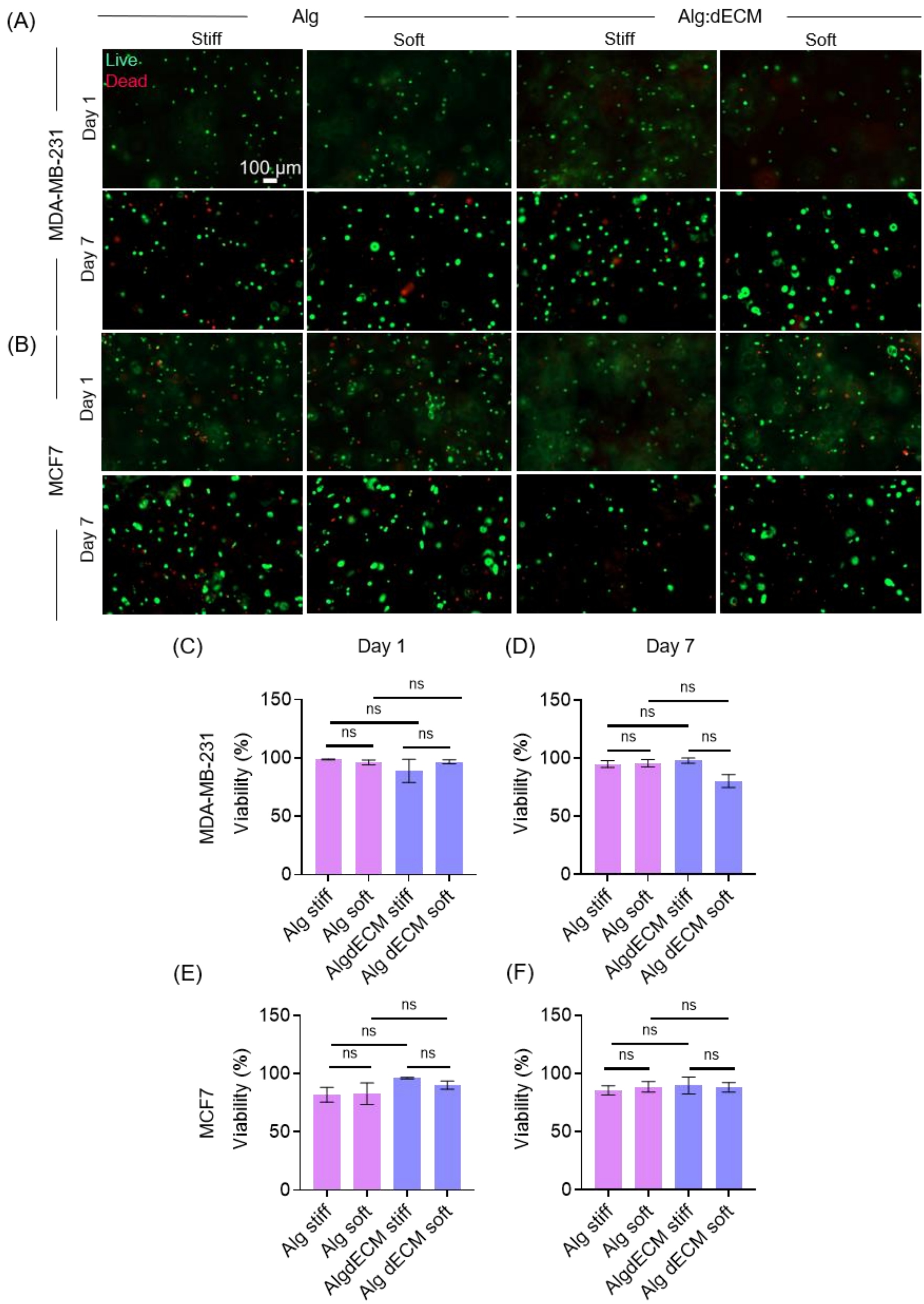


**Supplementary Figure S5**. Biocompatibility of Alg and Alg-dECM. A) Representative images of Z-stack projections of MDA-MB-231 and B) MCF7 cells stained with calcein-AM and ethidium homodimer-1 after 1 and 7 days of encapsulation, in the indicated high molecular weight 2.5 % w/v stiff (0.5 N:T) and soft (4.0 N:T)Alg and Alg-dECM hydrogels. C-D) Corresponding graphs depicting quantification of viability of MDA-MB-231 and E-F) MCF7 cells. n = 3 gels per conditions. Bar graphs display means with standard deviations. Mann-Whitney U test.

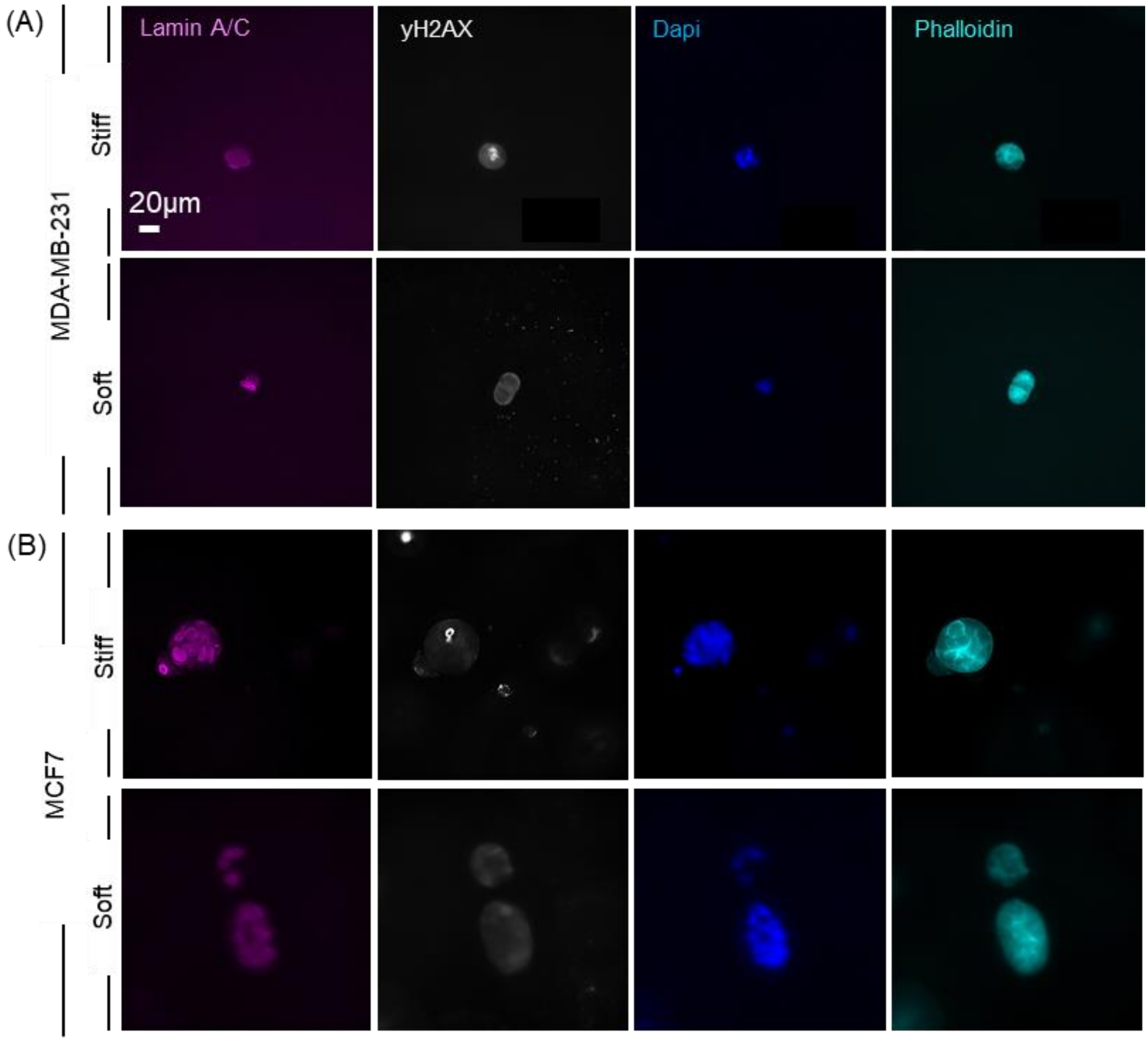


**Supplementary Figure S6**. Nuclear and genomic integrity in stiff and soft alginate. A) Representative images of Z-stack projections of MDA-MB-231 and B) MCF7 encapsulated in soft and stiff alginate stained with Lamin A/C (pink), yH2AX (white), Dapi (blue) and Phalloidin (turquoise).

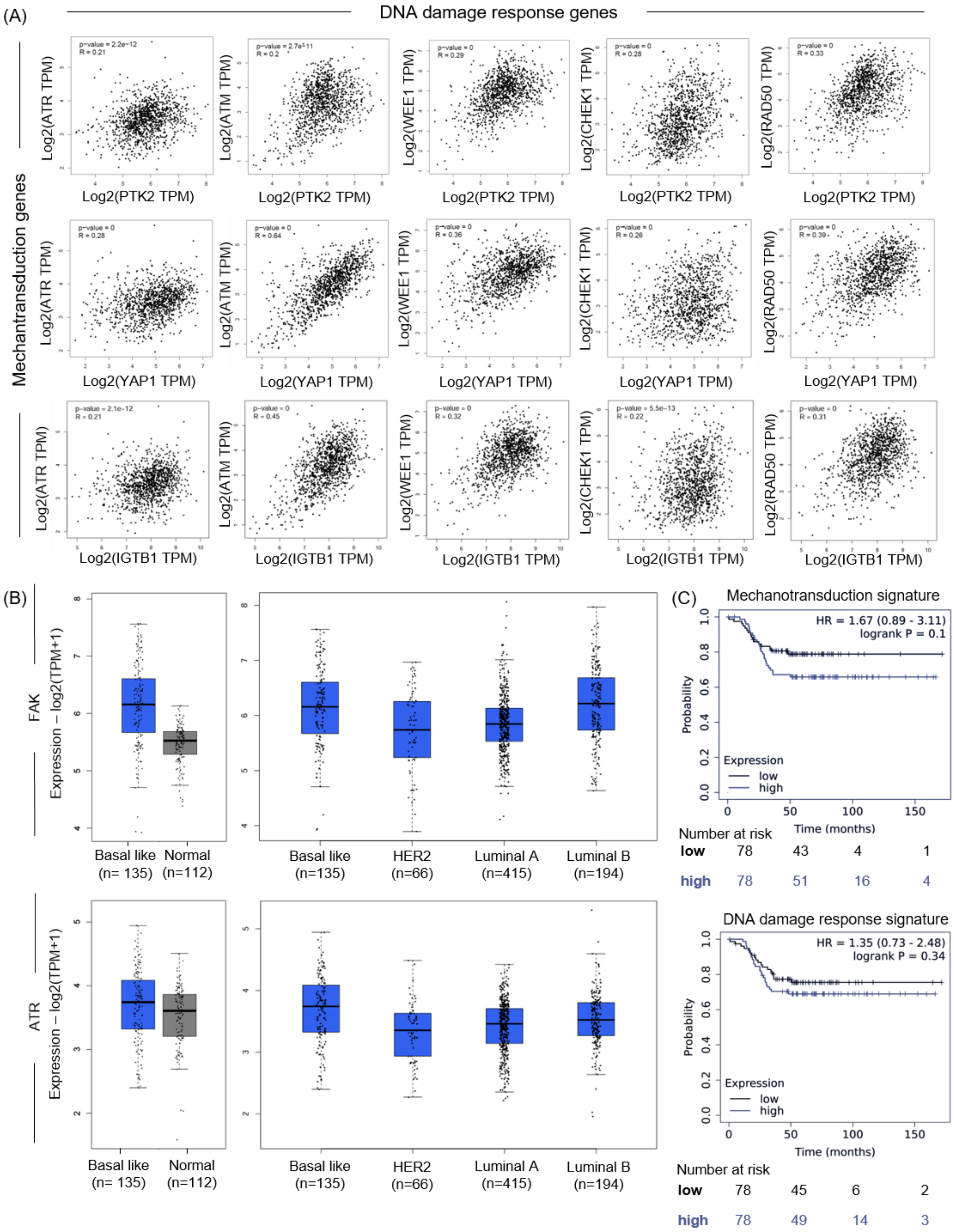


**Supplementary Figure S7.** Analysis of mechanotransduction and DNA damage in BC patients. A) Correlation analysis of DNA damage response genes with mechanotransduction genes (PTK2 = FAK gene) consistently showing positive associations. Data obtained from gepia2.cancer-pku.cn. [101] Expression of FAK and ATR in BC vs non-BC samples and across BC subtypes (n = number of patients). Data obtained from gepia2.cancer-pku.cn. [101] C)

Kaplan-Meier plot showing trend of worse distant metastasis free survival of TNBC patients expressing high mechanotransduction and DNA damage response signatures. Data obtained from kmplot.com. [102]

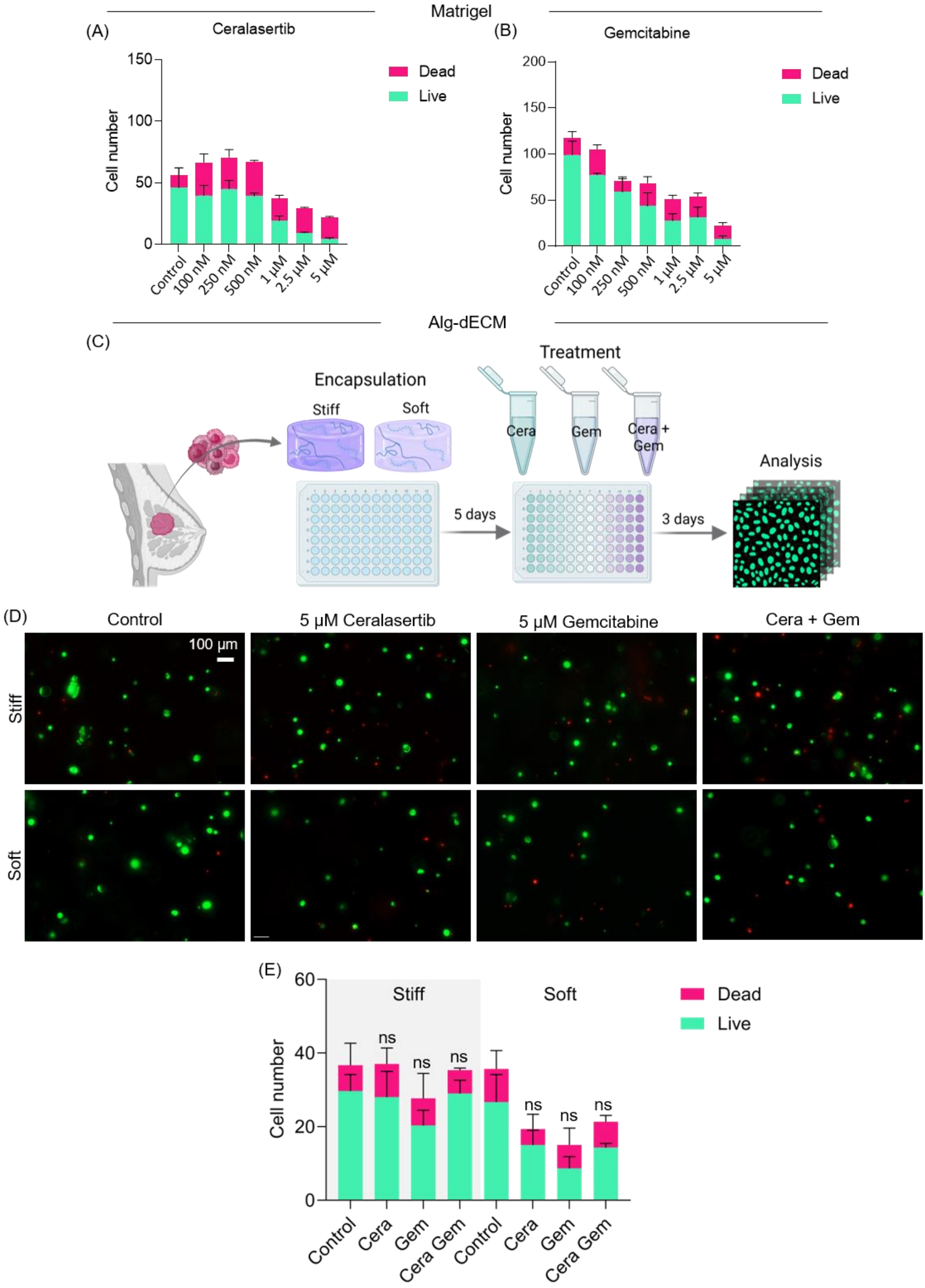


**Supplementary Figure S8**. Drug testing with patient-derived TNBC cells. A-B) Drug responses in Matrigel showing sensitivity in a concentration dependent manner to ATR inhibition via Ceralasertib (Cera) and chemotherapy with Gemcitabine (Gem). C) Schematic representation of drug testing with patient-derived TNBC cells in stiff and soft Alg-dECM. D) Representative images of Z-stack projections of patient-derived TNBC cells stained with calcein-AM and ethidium homodimer-1. E) Quantification of viability after 3 days of treatment with 5 µM Cera, 5 µM Gem or 5 µM Cera + 5 µM Gem. n = 3 gels per condition. Bar graphs show mean with standard deviation. Mann-Whitney U test.
